# The windblown: possible explanations for dinophyte DNA in forest soils

**DOI:** 10.1101/2020.08.07.242388

**Authors:** Marc Gottschling, Lucas Czech, Frédéric Mahé, Sina Adl, Micah Dunthorn

## Abstract

Dinophytes are widely distributed in marine- and fresh-waters, but have yet to be conclusively documented in terrestrial environments. Here we evaluated the presence of these protists from an environmental DNA metabarcoding dataset of Neotropical rainforest soils. Using a phylogenetic placement approach with a reference alignment and tree, we showed that the numerous sequencing reads that were assigned to the dinophytes did not associate with taxonomy, environmental preference, nutritional mode, or dormancy. All the dinophytes in the soils are most likely windblown dispersal units of aquatic species, and are not biologically active residents of terrestrial environments.

---

Environmental high-throughput sequencing (HTS) studies of protists have now been performed for over a decade (Santoferrara et al. 2020). During that time, a large diversity of dinophyte DNA sequences has also been uncovered. Dinophytes are an ecologically and economically important group of protists that exhibit many types of life styles and nutritional modes, including phototrophic, mixotrophic and heterotrophic forms as well as some being parasitic (Saldarriaga and Taylor 2017). All known dinophytes are from marine or freshwater environments (Adl et al. 2019). As they constitute a considerable fraction of the plankton and play an important role in the global aquatic ecosystem, HTS studies have detected dinophytes from waters sampled from the polar regions through to the tropics (de Vargas et al. 2015; Le Bescot et al. 2016; Elferink et al. 2017; Decelle et al. 2018; Lentendu et al. 2018; Annenkova et al. 2020; Giner et al. 2020; Gottschling et al. 2020). HTS studies have also detected DNA of dinophytes in terrestrial environments (Bates et al. 2013; Geisen et al. 2015; Mahé et al. 2017; Venter et al. 2017; Voss et al. 2019), although they are not expected to be there.

Aquatic protists can sometimes be detected in terrestrial environments, notably riparian soil, such as foraminifera (Meisterfeld et al. 2001; Lejzerowicz et al. 2010) and possibly haptophytes (Mahé et al. 2017). However, that does not mean the normally aquatic protists are biologically active in soils or other drier environments (Geisen et al. 2018). In the absence of observing putative soil dinophytes using direct microscopic observations, here we used Mahé et al.’s (2017) metabarcoding data from three lowland Neotropical rainforest soils to ask if the presence of dinophytes in those soils associate with taxonomy, environmental preference, nutritional mode, or dormancy.

## MATERIALS AND METHODS

### Environmental sampling and data

Sampling and sequencing of tropical soils originally took place in lowland rainforest in Costa Rica, Panama, and Ecuador (Mahé et al. 2017). The extracted soils DNAs were amplified for the hyper-variable V4 region of the SSU-rRNA locus using general eukaryotic primers (Stoeck et al. 2010); this short region has relatively strong phylogenetic signal, although it is not as strong as the full-length SSU-rRNA (Dunthorn et al. 2014; Gottschling et al. 2020). Illumina sequencing reads were clustered into OTUs using Swarm v2 (Mahé et al. 2015) and taxonomically assigned to the Protist Ribosomal Reference database (Guillou et al. 2013) using VSEARCH (Rognes et al. 2016). The 269 OTUs that were assigned to the dinophytes by Mahé et al. (2017), were extracted and used here for phylogenetic placements (**File S1**).

### Reference tree

From GenBank, 228 ingroup dinophytes, plus 10 outgroups, were downloaded, then aligned with MAFFT v6.624b (Katoh and Standley 2013) using the –auto option. Based on previous analyses (Gottschling et al. 2012, 2020; Žerdoner Čalasan et al. 2019), the full sequences of each species were used without excluding ambiguously aligned positions sites. Phylogenetic inferences of the reference alignment were carried out by using Maximum Likelihood (ML) as described in detail by Gottschling et al. (2012, 2020), using RAxML v8.2.10 (Stamatakis 2014) with the GTR+G substitution model. To determine the best fitted ML tree, we executed 10-tree searches from distinct random stepwise addition sequence Maximum Parsimony starting trees and performed 1,000 non-parametric bootstrap replicates. Reference alignment and tree available upon request.

### Phylogenetic placement of environmental OTUs

The OTU representative sequences obtained from Swarm were aligned against the reference alignment using PaPaRa v2.0 (Berger and Stamatakis 2011), and phylogenetically placed onto the ML reference tree using the Evolutionary Placement Algorithm (EPA) of RAxML. Next, all OTUs that were placed with at least 95% probability (combined likelihood weight ratios) in the dinophyte clade were extracted and visualized, using Gappa (Czech et al. 2020). For details on the extraction, see Czech et al. (2018); details of the workflow are published in a GitHub code repository (https://github.com/lczech/dinoflagellate-paper).

## RESULTS AND DISCUSSION

For the dinophyte reference alignment and tree, we included a broad representative taxon sample covering the known DNA sequence diversity with comprehensive sequence information. The alignment was 7,270 bp long and had 3,753 parsimony informative sites (52%, 15.7 per terminal taxon). The ML tree had many bipartitions that had high if not maximal bootstrap values. The Dinophyceae was inferred to be monophyletic, and it contained well-known subclades: Dinophysales, Gonyaulacales, Gymnodiniales, Peridiniales, Prorocentrales, †Suessiales as well as Amphidomataceae, Brachydiniaceae, and Tovelliaceae. Only 207 Neotropical soil OTUs from from the Mahé et al.’s (2017) that were assigned to the dinophytes, phylogenetically placed across the reference tree with high likelihood weight scores (**Fig. 1, File S2**).

**Figure 1:**
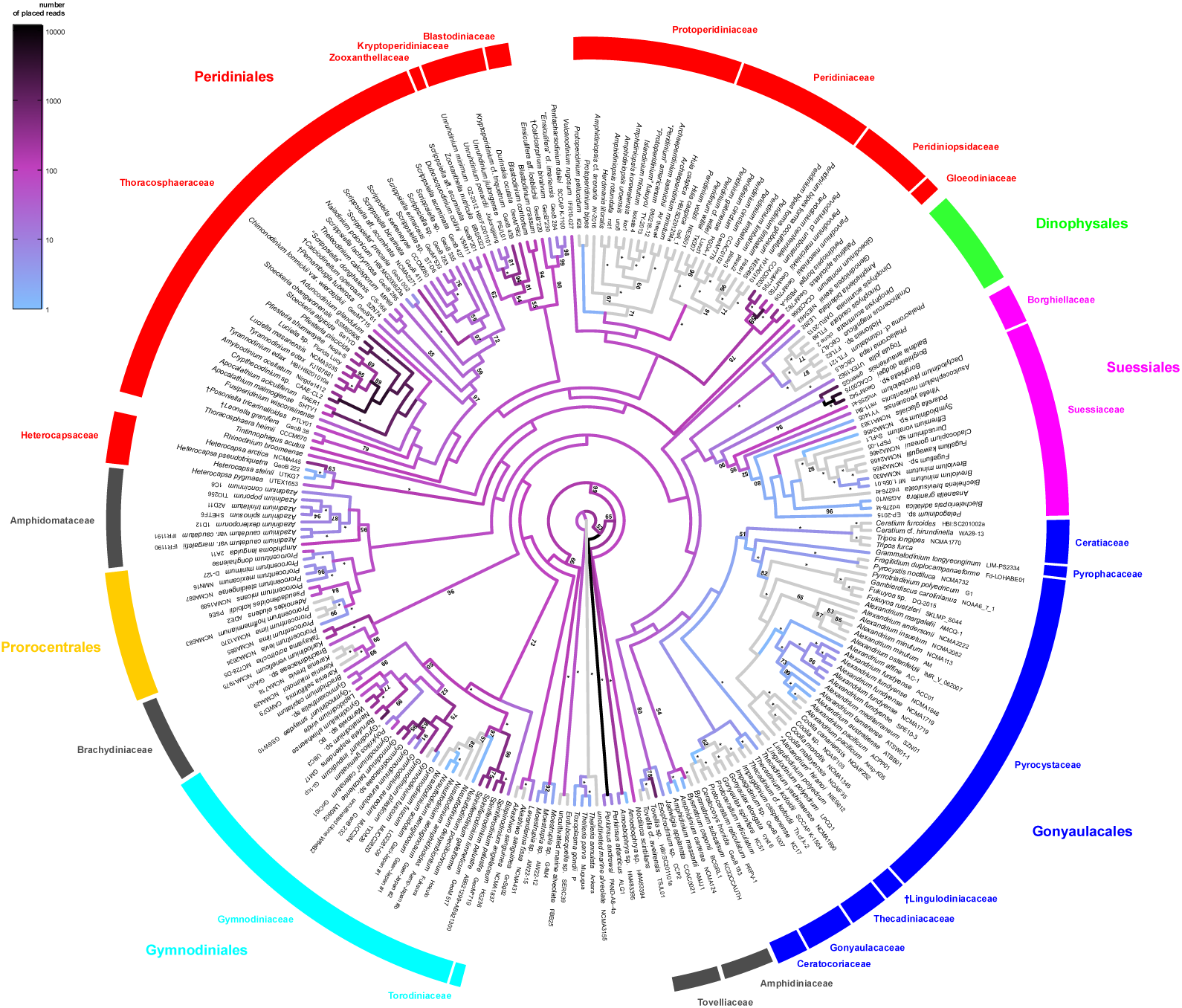
A molecular reference tree recognising major groups of dinophytes. Maximum likelihood (ML) tree of 228 representative dinophyte sequences (with strain number information), plus 10 outgroups, as inferred from a SSUrRNA nucleotide alignment (3,767 parsimony informative positions). Numbers on branches are ML bootstrap (above) and Bayesian support values (below) for the clusters (* = maximal support values; values <50 not shown).

There were no exclusive associations with taxonomy. Some of the dinophyte OTUs formed distinct clades of sequences that were unknown until the present study. However, the amount of such undescribed diversity is low compared to other microbial lineages such as the Fungi (Jones et al. 2011; Rosling et al. 2011). They placed onto early branches, which comprise heterotrophic, mainly parasitic species (Saldarriaga et al. 2003; Gómez et al. 2009; Bachvaroff et al. 2012; Gu et al. 2013), but also within the Peridiniales. Most of the OTUs, though, placed within already known lineages of the Gymnodiniaceae, Peridiniales and †Suessiales, a truly heterogeneous set of dinophytes including unarmored and thecate algae as well. There is no morphological trait that the OTUs would therefore necessarily share and that would unite them with these different taxa. Some thecate groups such as Dinophysales, Gonyaulacales, Prorocentrales and Protoperidiniaceae (Peridiniales) did not include any of the OTUs obtained from the environmental samples.

There were no exclusive associations with habitat preference. Some of the dinophyte OTUs placed onto branches that contain freshwater species, such as the Gymnodiniaceae (Kretschmann et al. 2015; Romeikat et al. 2020), Tovelliaceae (Lindberg et al. 2005), and peridinialean *Naiadinium* comprising a freshwater lineage within otherwise marine *Scrippsiella* s.l. (Kretschmann et al. 2014; Luo et al. 2016). This placement of OTUs within freshwater clades was similar to what was shown for the haptophyte OTUs from the same rainforest soils (Mahé et al. 2017). However, none of the OTUs placed with the Peridiniaceae, which is one of the most prominent freshwater dinophyte lineages (Moestrup and Calado 2018), but some OTUs placed on branches that contain just marine species, such as the Amphidomataceae (Tillmann et al. 2014) and Brachydiniaceae (Bergholtz et al. 2006; Henrichs et al. 2011). Groups such as Dinophysales, Gonyaulacales, Prorocentrales and Protoperidiniaceae (Peridiniales) are primarily marine and did not include any of the OTUs obtained from the environmental samples.

There were no exclusive associations with nutritional mode. Some of the dinophyte OTUs placed onto branches that predominantly contain phototrophic species, such as Gymnodiniaceae, Peridiniales and †Suessiales. Other OTUs placed with the heterotrophic species such as in Thoracosphaeraceae (i.e., *Pfiesteria* and relatives), but not in the consistently heterotrophic Dinophysales and Protoperidiniaceae (Peridiniales). Transitions between phototrophic and heterotrophic modes are thought to occur in the dinophytes (Jeong et al. 2012; Fawcett and Parrow 2014), but there is no phylogenetic signal for this trait at high taxonomic levels. Additionally, some OTUs placed onto early branches that include many parasitic species (Saldarriaga et al. 2003; Gómez et al. 2009; Bachvaroff et al. 2012; Gu et al. 2013), and they placed onto younger branches that also include parasites, such as the Gymnodiniaceae (Gómez et al. 2009; Kretschmann et al. 2015; Romeikat et al. 2020) and Peridiniales (Coats et al. 2010; Gottschling and Söhner 2013).

There were no exclusive associations with dormancy practice. In addition to flagellated trophic cells, coccoid stages are integral part in the life-history of many dinophytes from marine and freshwater environments. Coccoid cells are particularly abundant in the Gonyaulacales where no OTUs placed, or in the Peridiniales where many OTUs placed (Dale 1983; Evitt and Davidson 1964; Fensome et al. 1993; Wall 1971). Exact functions of coccoid cells are not worked out rigorously for more than a handful of dinophyte species, but may frequently correspond to resting and/or dormancy stages (Fensome et al. 1993). Deposited in sediments, coccoid cells have the potential to preserve the local biodiversity like diaspores in a seed bank (Dale 1983). They can be considered either as old remnants from former aquatic and today terrestrial habitats (Boere et al. 2011; Sønstebø et al. 2010) or the result of random dispersal (Foissner 2006, 2011) and subsequent loss in terrestrial habitats. However, the difference between the taxonomic assignments of the rainforest soil samples (see above) may indicate that the ability to form coccoid cells during life-history is not decisive for their terrestrial occurrence. An ecological group that has been receiving more interest in the past years and that may also be considered for the evaluation of the terrestrial samples are benthic dinophytes living in the intertidal (Hoppenrath et al. 2014); phylogenetically, it is a heterogeneous assemblage recruiting members particularly from Gonyaulacales and Peridiniales. But the bigger question remains, as to whether there are any soil dinophytes at all, or we are simply detecting windblown cells or dormant cells.

## Conclusion

The presence of dinophyte DNA sequences in the Neotropical rainforest soils—as an exemplar of a terrestrial environment—did not associate with taxonomy, environmental preference, dormancy practice during life history, or nutrition mode/organismal interaction. The reason for the presence thus remains to be identified. We have to keep in mind that presence of DNA does not necessarily indicate biological activity. The environmental DNA sequences identified here are scattered unevenly across the classification, but it may represent more easily dispersed surface algae. To completely resolve this paradox, microscopy of soil samples with dinophyte DNA needs to be performed to verify their presence; however, in two centuries of looking at soil samples, dinophytes have never been observed. The most likely explanation for dinophyte DNA in forest soils is that they were passively dispersed there— they are the windblown.

## ACKNOWLEDGMENTS

We thank Alexandros Stamatakis for computational support. Funding came from the Deutsche Forschungsgemeinschaft (grant DU1319/5-1) to MD.

