## Supplementary material for "The windblown: possible explanations for dinophyte DNA in forest soils": File S1

>d6a793bd001362ed30a1ba5f8a06fa8dd60561c7_19311
agctccaatagcgtatattaaagttgttgcggttaaaaagctcgtagttggacttctgcgaaggacgaccggtccgcccaccgggtgagtatctggttcggcctttgcatcctcttggagaaggtgtgttcacttcactgtgtgcacccgtatccaggacattttactttgaggaaattggagtgcttcaagcaggcacacgccttgaacacatgagcatggaataatgcgataggacctcggttctattttgttggtttctagcactgaggtaatgataaatagggacagttgggggcattcgtatttaactgtcagaggtgaaattcttggatttgttaaagacggaccactgcgaaagcatttgccacggatgtcttca
>4c37ff12e573dedee33c48df1de19af4033e5784_11879
agctccaatagcgtatattaaagttgttgcagttaaaaagctcgtagttgaatttctgctggagttgaataggtctgctctttgagtgtgtaccttgtttcgacttcgcatcctcttagaatgttattaaagtactttattgtgctttatataatatctaagatctttactttgagaaaattagagtgtttcaagcaggcatattgccttgaatactttagcatggaataataaaataggattttggttctattttattggtttatagaactaaagtaatgattaatagggacagttgggggcattcgaatttaactgtcagaggtgaaattcttagatttgttaaagtcgaactactgcgaaagcatttgccaaggatgttttca
>6d961ec230c74df89d504c2908ba347446d9071d_11712
agctccaaaagcgtatattaaagttgttgcggttaaaaagctcgcagttggattttatagtcggagctacattttggtgtaactcgattatggtaataatttattattacacgttactttgaggaaattagagtgtttaaagcagatctttgtcatgtatatattagcatggaataacactaaggattattaaaactttgttggttactttttttaataatgattgatagggacagccgggggcatttgtatttaacagtcagaggtggaattcttggatttgttaaggacaaactattgcgaaagcatttgccaaggatgttttcg
>b8e2af4b0ca8de85d4643140f299e0a6a5eb1e51_17104
agctccaatagcgtatattaaagttgttgcggttaaaaagctcgtagttggatttctgcttaggacgaccggtccgcccactgggtgagtatctggctcggcctgggcatcttcttggagaacgtagctgcgcttgattgtgtggtgcggaatccaggacttttactttgagaaaattagagtgtttcaagcaggcacacgccttgaatacattagcatggaataataatataggacctcggttctattttgttggtttctagagctgaggtaatgattgatagggatagttgggggcattcgtatttaactgtcagaggtgaaattcttggatttgttaaagacggactactgcgaaagcatttgccaaggatgttttca
>d20c9dfc8a9becdb9cda48bd9365c16b68205322_7565
agctccaaaagcgtatattaaagttgttgcggttaaaaagctcgtagttggattttatagtcgaagctacattttggtgtaactcgattatggtaataatttattattacacgttactttgaggaaattagagtgtttaaagcagatctttgtcatgtatatattagcatggaataacactaaggattattaaaactttgttggttactttttttaataatgattgatagggacagccgggggcatttgtatttaacagtcagaggtggaattcttggatttgttaaggacaaactattgcgaaagcatttgccaaggatgttttcg
>ee53ac3750359fda9d8a011237943062f9c3d66c_5772
agctccaatagcgtatattaaagttgttgcagttaaaaagctcgtagttggatttctgccttcgatgggtgccccaccttagcggtgcgtgacgttcaatcgctggcattccttcactgtgcgaaagcacacgtgatactttactttgagtaaattagagtgtttcaagcaggcgtttgcctgaatactacagcatggaataatgagataggacttctgttctttctttattggttctagggatggaagtaatggttaatagggacagtcgggggcattcgtattcaactgttagaggtgaaattcttggattagttgaagaccaacaactgcgaaagcatttgccaaggatgttccca
>4c554190d2d6dd436de4e6c438462f619e18265f_5365
agctccaatagcgtatattaaagttgttgcggttaaaaagctcgtagttggatttctgttgaggacgatcggtccgccttctgggtgagtatctggctcggccttggcatcttcttggagaacgtactgcacttcattgtgtggtgcggaatccaggacctttactttgaggaaattagagtgtttcaagcaggcgcacgccttgaatacattagcatggaataataagataggacctcggttctattttgttggtttctagagctgaggtaatgattaatagggatagttgggggcattcgtatttaactgtcagaggtgaaattcttggatttgttaaagacggactactgcgaaagcatttgccaaggatgttttca
>fc96e6a67377f73fe72f4d4638fbec6c59194dc3_4520
agctccaatagcgtatattaaagttgttgcggttaaaaagctcgtagttggatttctgccaaggatgaccagtccgccttcgggtgtgtatctgtgttgaccgaggcatcttcctaggaaacgtctgtacttaactgtgcggcacgtattctaggacttttactttgaggaaattagagtgtttcaagcaggcatatgccttgaatacattagcatggaataataaggtaggactccggttctattttgttggtttctagaacaggagtaatgattaatagggacagtcgggggcattcgtactcaactgtcagaggtgaaattcttggatttgttggagacgaactactgcgaaagcatttgccaaggatgttttca
>7ee5cf32c634bd73780636ced48ccf80b5484854_4826
agctccaatagcgtatattaaagttgttgcggttaaaaagctcgtagttggatttctgtcgaggacgaccggtccgccctccgggtgagtatctggttcggcctaggcatcctcttggagaaggagcggtcactttgctgtgaacgcccgtatccaggacttttactttgaggaaattggagtgcttcaagcaggcacacgccttgaacacatgagcatggaataatgcgataggacctcggttctattttgttggtttctagagctgaggtaatgataaatagggacagttgggggcattcgtatttaactgtcagaggtgaaattcttggatttgttaaagacggaccactgcgaaagcatttgccacggatgtcttca
>532b5fb664c9995b8f0f58a1be8a1d6c27db6ebd_3064
agctccaatagcgtatattaaagttgttgcagttaaaaagctcgtagttgaatttctgctggagttgaataggtctgctctttgagtgtgtaccttgtttcgacttcgcatcctcttagaatgttattaaagtactttattgtgctttatataatatctaagatttttactttgagtaaattagagtgtttcaagcaggcatattgccttgaatactttagcatggaataataaaataggattttggttctattttattggtttatagaactaaagtaatgattaatagggacagttgggggcattcgaatttaactgtcagaggtgaaattcttagatttgttaaagtcgaactactgcgaaagcatttgccaaggatgttttca
>f9e3bb9f1cc7d1d39480a29bb220c52bc9b27608_2933
agctccaatagcgtatatttaagttgttgcagttaaaaagctcgtagttggatttctgccgaggacgaccggtccgcccactgggtgtgtatctggctcggcctgggcatcttcttggagaacgtagctgcacttgactgtgtggtgcggtatccaggacttttactttgaggaaattagagtgtttcaagcaggcacacgccttgaatacattagcatggaataataagataggaccttggttctattttgttggtttctagaactgaggtaatgattaatagggatagttgggggcattcgtatttaactgtcagaggtgaaattcttggatttgttaaagacggactactgcgaaagcatttgccaaggatgttttca
>93a188b7435e9ac5053755f2989f2cca340dba24_3263
agctccaatagcgtatattaaagttgttgcggttaaaaagctcgtagttggatttctgctaaggacgaccggtccgccttcgggtgtgtacatgtgttgaccgaggcatcaatctggaagacgcctgtacttaaccgtgcgggcgtaattcagacgttttactttgaggaaattagagtgtttcaagcaggcaattgccctgaatacattagcatggaataataagataggacttcggttctattttgttggtttctagaactgaagtaatgattaatagggacagtcgggggcattcgtactcaactgtcagaggtgaaattcttggatttgttgaagacgaactactgcgaaagcatttgccaaggatgttttca
>402ff1c02cf1114cb6b6cfe542997b9d276dedbd_2668
agctccaatagcgtatattaaagttgttgcggttaaaaagctcgtagttggatttctgccaaggacgaccggtccaccttagggtgtgtatctgtgttgaccgaggcatctttctaggtgacgcccgtgcttcactgtgcgacgcgtgctctagaacttttactttgaggaaattagagtgttccaagcaggcttatgccctgaatacattagcatggaataataaggtaggacttcggttctattttgttggtttctagaactgaagtaatgattaatagggacagtcgggggcattcgtactcaactgtcagaggtgaaattcttggatttgttggagacgaactactgcgaaagcatttgccaaggatgttttca
>92dac336af3373c32854d83a7535c7232bb2e264_2893
agctccaatagcgtatattaaagttgttgcggttaaaaagctcgtagttggatttctgcttaggatgaccggtccgccttcgggtgtgtatcagcgttgtccagagcatctttctagttgcgcccgtgcttcaatgtacggtgcgtgttctagatcttttactttgaggaaattagagtgtttcaagcaggcatatgccttgaatactttagcatggaataataagataggacttcggttctattttgttggtttctagaactgaagtaatgattaatagggacaatcgggggcattcgtactcaacagtcagaggtgaaattcttggatttgttggagacgaactactgcgaaagcatttgccaaggatgttttca
>e26590a7c401a8e17ddf3090947520f9f3c1fea3_2967
agctccaatagcgtatattaaatttgttgcggttaaaaagctcgtagttggatttctgccgaggacgaccggtccgcccactgggtgtgtatctggctcggcctgggcatcttcttggagaacgtagctgcacttgactgtgtggtgcggtatccaggacttttactttgaggaaattagagtgtttcaagcaggcacacgccttgaatacattagcatggaataataagataggaccttggttctattttgttggtttctagaactgaggtaatgattaatagggatagttgggggcattcgtatttaactgtcagaggtgaaattcttggatttgttaaagacggactactgcgaaagcatttgccaaggatgttttca
>d1f1e1ed2c659894415f7e0408f5dae1d6ecad2c_1593
agctccaatagcgtatattaaagttgttgcggttaaaaagctcgtagttggatttctgctgaggacgaccggtccgccctctgggtgagtatctggctcggccttggcatcttcttggagaacgttactgcacttgattgtgtggtgcggtatccaggacttttactttgaggaaattagagtgtttcaagcaggcacacgccttgaatacattagcatggaataataagataggaccttggttctattttgttggtttctagaactgaggtaatgattaatagggatagttgggggcattcgtatttaactgtcagaggtgaaattcttggatttgttaaagacggactactgcgaaagcatttgccaaggatgttttca
>f5390614fe3126281636dd5b8fa4cffaf8c3b26c_1991
agctccaatagcgtatattaaagttgttgcggttaaaaagctcgtagttggatttctgttgaggacgaccggtccgccctctgggtgagtatctggctcggccttggcatcttcttggggaacgttactgcacttgactgtgtggtgcggtatccaggacttttactttgaggaaattagagtgtttcaagcaggcgcacgccttgaatacattagcatggaataataagataggacctcggttctattttgttggtttctagagctgaggtaatgattaatagggatagttgggggcattcgtatttaactgtcagaggtgaaattcttggatttgttaaagacggactactgcgaaagcatttgccaaggatgttttca
>f5d50cc840e1144e93eeaa6d29b067d9ac79902e_1764
agctccaatagcgtatattaaagttgttgcggttaaaaagctcgtagttggatttctgctgaggacgaccggtccgccctccgggtgagtatctggttcggcctgagcatcctcttggagaaggtgcgtacacttcgttgtgcacgcccgtatccaggacttttactttgaggaaattggagtgcttcaagcaggcacacgccttgaacacatgagcatggaataatgcgataggacctcggttctattttgttggtttctagagctgaggtaatgataaatagggacagttgggggcattcgtatttaactgtcagaggtgaaattcttggatttgttaaagacggaccactgcgaaagcatttgccacggatgtcttca
>720928ac158e8a47cbdb1c68d0bb65aed01439e5_1788
agctccaatagcgtatattaaagttgttgcggttaaaaagctcgtagttggatttctgccaaggatgaccggcccgccttcgggtgtgtgcatgtgttgaccgaggcatttttctgatattggcttcacttcactgtgttgtcaatgttcagagatgttactttgaggaaattagagtgtttcaagcaggcaattgccctgaatacattagcatggaataacaatataggactccggttctattttgttggtttcttgagctggagtaatgattaatagggacaatcgggggcatccgtactcaaccgtcagaggtgaaattcttggattggttgaagacgaactactgcgaaagcatttgccaaggatgttttca
>793ed5f0b15e46106a691ff2ff3bb4ca11033d3e_2838
agctccaatagcgtatattaaagttgttgcggttaaaaagctcgtagttggatttctgccgaggacgaccggtccgcccactgggtgtgtatctggctcggcctgggcatcttcttggagaacgtatctgcacttgactgtgtggtgcggtatccaggacttttactttgaggaaattagagtgtttcaagcaggcacacgccttgaatacattagcatggaataataagataggaccttggttctattttgttggtttctagagctgaggtaatgattaatagggatagttgggggcattcgtatttaactgtcagaggtgaaattcttggatttgttaaagacggactactgcgaaagcatttgccaaggatgttttca
>5e197728f5006b8e36139eabb3d4562d4c5df937_1488
agctccaatagcgtatattaaagttgttgcggttaaaaagctcgtagttggatttctgctgaggacgaccggtccgacctctgggtgagtatctggatcggcctaggcatcttcttggagaacgtagctgcacttgactgtgtggtgcggaatccaggacttttactttgagaaaattagagtgtttcaagcaggcacacgccttgaatacattagcatggaataatacttcaagaccttggttctattttgttggtttctagatccagggtaatgattgatagggatagttgggggcattcgtatttaactgtcagaggtgaaattcttggatttgttaaagacggactactgcgaaagcatttgccaaggatgttttca
>9026474cc46a75f00939957b1a6a708f879fba22_922
agctccaatagcgtatattaaagttgttgcagttaaaaagctcgtagttgaatttctgctggagttgaataggtctgctctttgagtgtgtacctggtttcgacttggcatcctcttagaatgttattaaagtactttattgtgctttatataatatctaagatctttactttgagtaaattagagtgtttaaagcaggcatattgccttgaatactttagcatggaataataaaataggactttggttctattttattggtttatagaactaaagtaatgattaatagggacagttgggggcattcgaatttaactgtcagaggtgaaattcttagatttgttaaagtcgaactactgcgaaagcatttgccaaggatgttttca
>08e76210f82f3c85a0795393e744a2b37a2be132_986
agctccaaaagcgtatattaaagttgttgcggttaaaaagctcgtagttggattttatagtcgaagctattttggtgtaactcgattatggtaataatttattattacacgttactttgaggaaattagagtgtttaaagcagatctttgtcatgtatatattagcatggaataacactaaggattattaaaactttgttggttactttttttaataatgattgatagggacagccgggggcatttgtatttaacagtcagaggtggaattcttggatttgttaaggacaaactattgcgaaagcatttgccaaggatgttttcg
>6698fbde9ac45aba6fab523c102479eb1fa4988f_953
agctccaatagcgtatattaaagttgttgcggttaaaaagctcgtagttggatttctgctgaggaagaccggtctgccctctgggtgagtatctggctccgccttggcatcttcttgggaaccgttgctgcacttgattgtgtggtgcgggattcaggacatttactttgaggaaattagagtgcttcaagcaggcatggtgtctgaatacattagcatggaataataagatgggccctggctctgttttgttggtttctggagccgaggtcatgattaacagggacaattgggggcattcgtacttaactgtcagaggtgaaattcttggatttgttaatgacggactactgcgaaagcatttgccaaggatgttttca
>4d66b77aead4818a5d3d13f9f27ee17d2164f1d7_987
agctccaatagcgtatattaaagttgttgcggttaaaaagctcgtagttggatttctgctgaggacgaccggtccgccctctgggtgagcatctggctcggcctgagcatcctcttggagaaggtgtgtgcactttactgtgtgcacccgtatccaggacttttactttgaggaaattggagtgcttcaagcaggcacacgccttgaacacatgagcatggaataatgcgataggacctcggttctattttgttggtttctagaactgaggtaatgataaatagggacagttgggggcattcgtatttaactgtcagaggtgaaattcttggatttgttaaagacggaccactgcgaaagcatttgccacggatgtcttca
>b30350baadcc2d1e6e13db7ac9b5aadb37207dd9_864
agctccaatagcgtatattaaagttgttgcagttaaaaagctcgtagttgaatttctgcgagagttgaataggtctgctctttgagtgtgtaccttatttcgacttcgcatcctcttagaatgttataaaagtactttattgtgctttttataatatctaagatttttactttgagtaaattagagtgtttcaagcaggcatattgccttgaatactttagcatggaataataagataggattttggttctattttgttggtttatagaactaaagtaatgattaatagggacagttgggggcattcgaatttaactgtcagaggtgaaattcttagatttgttaaagtcgaactactgcgaaagcatttgccaaggatgttttca
>aba964e993016c42c28a49f572a9176a7424e41e_913
agctccaatagcgtatattaaagttgttgcggttaaaaagctcgtagttggatttctgctgaggacgaccggtctgcccaccgggtatgcatctggatcggcctgagcatcttcttggcgaacgctgctgcactttactgtgtggtgcggtatccaggacttttactttgaggaaattagagtgcttaaagcaggcatacgcctgaacacatgagcatggaataatatgataggacctcggttctattttgttggtttctagaactgaggtaatgataaatagggacagttgggggcattcgtatttaactgtcagaggtgaaattcttggatttgttaaagacggaccactgcgaaagcatttgccacggatgtcttca
>159192736cbb97e1d1079f0709fd7841dadc1d19_866
agctccaaaagcgtatattaaagttgttacggttaaaaagctcgtagttggattttatagtcgaagctacattttggtgtaactcgattatggtaataatttattattacacgttactttgaggaaattagagtgtttaaagcagatctttgtcatgtatatattagaatggaataacactaaggattattaaaactttgttggttactttttttaataatgattgatagggacagccgggggcatttgtatttaacagtcagaggtggaattcttggatttgttaaggacaaactattgcgaaagcatttgccaaggatgtttttg
>dc3a7370b8e168d66c6d83d512081a55524571e0_1176
agctccaatagcgtatattaaagttgttgcagttaaaaagctcgtagttgaatttctgctggagttgattaggtctgctctgtgagtgtgtaccttaatatcgacttagcatcctcttagaatgttattaaagtactttattgtgctttatataatatctaagatttttactttgagtaaattagagtgtttatagcaggcatattgccttgaatactttagcatggaataataaaataggattttggttctattttattggtttatagaactaaagtaatgattaatagggacagttgggggcattcgaatttaactgtcagaggtgaaattcttagatttgttaaagtcgaactactgcgaaagcatttgccaaggatgttttca
>4927cf57794970028dd04aa6cbd89a4937834072_566
agctccaatagcgtatattaaagttgttgcggttaaaaagctcgtagttggatttctgcttaggacgaccggtccgcccactgggtgagtatctggctcggcctgggcatcttcttggagaacgtagctgcgcttgattgtgtggtgcggaatccaggacttttactttgagaaacttagagtgtttcaagcaggcacacgccttgaatacattagcatggaataataatataggacctcggttctattttgttggtttctagagctgaggtaatgattgatagggatagttgggggcatttgtatttaactgtcagaggtgaaattcttggatttgttaaagacggactactgcgaaagcatttgccaaggatgttttca
>e2f98c7514e8a82aff090282cece162308a948c4_1246
agctccaatagcgtatattaaaattgttgcagttaaaaagctcgtagttgaatttctgctggagttgaataggtctgctctttgagtgtgtaccttgtttcgacttcgcatcctcttagaatgttattaaagtactttattgtgctttatataatatctaagatttttactttgagtaaattagagtgtttaaagcaggcatattgccttgaatactttagcatggaataataaaataggactttggttctattttattggtttatagaactaaagtaatgattaatagggacagttgggggcattcgaatttaactgtcagaggtgaaattcttagatttgttaaagtcgaactactgcgaaagcatttgccaaggatgttttca
>9d9b7584bef62b3c85d708da59bb2f6556d5efde_505
agctccaatagcgtatattaaagttgttgcggttaaaaagctcgtagttggatttctgccgaggacgaccggtccgcccactgggtgtgtatctggctcggcctgggcatcttcttggagaacgttactgcacttgattgtgtggtgcggtatccaggacttttactttgaggaaattagagtgtttcaagcaggcgcacgccttgaatacattagcatggaataataagataggacctcggttctattttgttggtttctagagctgaggtaatgattgatagggatagttgggggcattcgtatttaactgtcagaggtgaaattcttggatttgttaaagacggactactgcgaaagcatttgccaaggatgttttca
>1ce825f5402141d6410f1766f3916e5c6d943c59_840
agctccaatagcgtatattaaagttgttgcggttaaaaagctcgtagttggatttctgccgaggacgaccggtccgcccactgggtgtgtatctggctcggcctgggcatcttcttggaggacgtatctgcacttgactgtgtggtgcggcatccaggacttttactttgaggaaattagagtgtttcaagcaggcacacgccttgaatacattagcatggaataataaggtaggaccttggttctattttgttggtttctagaactgaggtgatgattaatagggatagttgggggcattcgaatttaactgtcagaggtgaaattcttggatttgttaaagacgaactactgcgaaagcatttgccaaggatgttttcg
>a0fd5c3584dad6151533e2e5545ecd105c622188_388
agctccaatagcgtatattaaagttgctgcagttaaaaagctcgtagttggatttctgcttaggacgaccggtccgcccactgggtgagtatctggctcggcctgggcatcttcttggagaacgtagctgcgcttgattgtgtggtgcggaatccaggacttttactttgagaaaattagagtgtttcaagcaggcacacgccttgaatacattagcatggaataataatataggacctcggttctattttgttggtttctagagctgaggtaatgattgatagggatagctgggggcattcgtatttaactgtcagaggtgaaattcttggatttgttaaagacggactactgcgaaagcatttgccaaggatgttttca
>e1e8bdf388cb28c352b31143811a618cee79ef77_373
agctccaatagcgtatattaaagttgttgcggttaaaaagctcgtagttggatttctgccgaggacgaccggtccgccctctgggtgagtatctggctcggcctgggcatcttcttggagaacgtatctgcacttgactgtgtggtgcggtattcaggacttttactttgaggaaattagagtgtttcaagcaggcatacgccttgaatacattagcatggaataataagataggacctcggttctattttgttggtttctagatctgaggtaatggttaatagggatagttgggggcattcgtatttaactgtcagaggtgaaattcttggatttgttaaagacggactactgcgaaagcatttgccaaggatgttttca
>0e13994ab91bce37f31dad5fd0949ed600a4b434_360
agctccaatagcgtatattaaagttgttgcagttaaaaagctcgtagttggatttctgcttaggacgaccggtccgcccactgggtgagtatctggctcggcctgggcatcttcttggagaacgtagctgcgcttgattgtgtggggcggaatccaggacttttactttgagaaaattagagtgtttcaagcaggcacacgccttgaatacattagcatggaataataatataggacctcggttctattttgttggtttctagagctgaggtaatgattgatagggatagttgggggcattcgtatttaactgtcagaggtgaaattattggatttgttaaagacggactactgcgaaagcatttgccaaggatgttttca
>c26186e453d2655a42433bc36cf20b587a21a457_526
agctccaatagcgtatattaaagttgttgcggttaaaaagctcgtagttggatttctgttgaggacgatcggtccgccttctgggtgagtatctggctcggccttggcatcttcttggagaacgtactgcacttcattgtgtggtgcggaatccaggacctttactttgaggaaattagagtgtttcaagcaggcgcacgccttgaatacattagcatggaataataagataggacctcggttctatttaatagggatagttgggggcattcgtagttaactgtcagaggtgaaattcttggatttgttaaagacggactactgcgaaagcatttgccaaggatgttttca
>856a5e57aeaf7fbc8e554ef054a7f8d1dc1c7419_587
agctccaatagcgtatatttaagttgttgcagttaaaaagctcgtagttggatttctgcttaggacgaccggtccgcccactgggtgagtatctggctcggcctgggcatcttcttggagaacgtagctgcgcttgattgtgtggtgcggaatccaggacttttactttgagaaaattagagtgtttcaagcaggcacacgccttgaatacattagcatggaataataatataggacctcggttctattttgttggtttctagagctgaggtaatgattgatagggatagttgggggcattcgtatttaactgtcagaggtgaaattcttggatttgttaaagacggactactgcgaaagcatttgccaaggatgttttca
>8deba22c1e1d6883a2b33a0bf42c989cbd5fca23_643
agctccaatagcgtatattaaagttgttgcggttaaaaagctcgtagttagatttttgatctagtcgaccggtcactccaatggaatgtatcaggtttgattagaacatttgtctgaaatttttgtctgcacttcactgtgtggtcaaagattcagaccttttactttgaggaaattagagtgtttcaagcaggcatttgccttgaatacgttagcatggaataataaaataagactttggttttattttgttggtttctaaaactaaagtaatgattaatagggatagtcgggggcattcgtacttaactgtcagaggtgaaattcttggatttgttaaagacgaactactgcgaaagcatttgccaaggatgttttca
>78d3f5bf16caf870fd8b2510c52ae4d38b48e8fe_359
agctccacaagtgtatattaaaattgttgcggttgaagagttcgtagttgttcgttcgaattccaggcttatggaatttgacagggcttgacagccttttgctttggataaacaaaagtgttcaatttagcttttgcggattttggtctgcagtacaattactttactgccaacgtgatggttggtagttcatcctttaggaaggcgcaaagggatattggtttcttgcgaacagaggtgaaattcttggactcgcaagcgaccagcagctgcgaaggtgtttatcaagggcgcgcttc
>98a6d25c852f97c65fca4e09121b3f88756272f9_310
agctccaatagcgtatattaaagttgttgcagttaaaaagctcgtagttgaatttctgcgagagttgaataggtctgctctttgagtgtgtaccttatttcgacttcgcatcctcttagaatgctataaaagtactttattgtgctttttataaaatctaagatttttactttgagtaaattagagtgtttcaagcaggcatattgccttgaatactttagcatggaataataagataggactttggttctattttgttggtttatagaactaaagtaatgattaatagggacagttgggggcattcgaatttaactgtcagaggtgaaattcttagatttgttaaagtcgaactactgcgaaagcatttgccaaggatgttttca
>da05ba49bf3abdc15426a10f9f4b399e868fc1b0_317
agctccaatagcgtatattaaagttgttgcagttaaaaagctcgtagttgaatttctgcgagagttgaataggtctgctctttgagtgtgtaccttatttcgacttcgcatcctcttagaatgttataaaagtactttattgtgctttttataatatctaagatctttactttgagtaaattagagtgtttcaagcaggcatattgccttgaatactttagcatggaataataagataggattttggttctattttattggtttatagaactaaagtaatgattaatagggacagttgggggcattcgaatttaactgtcagaggtgaaattcttagatttgttaaagtcgaactactgcgaaagcatttgccaaggatgttttca
>ee99783001b9b4c561ea052f5355a01588c49ddb_207
agctccaatagcgtatattcaagttgttgcggttaaaaagctcgtagttggatttctgcttaggacgaccggtccgcccactgggtgagtatctggctcggcctgggcatcttcttggagaacgtagctgcgcttgattgtgtggtgcggaatccaggacttttactttgagaaaattagagtgtttcaagcaggcacacgccttgaatacattagcatggaataataatataggacctcggttctattttgttggtttctagagctgatgtaatgattgatagggatagttgggggcattcgtatttaactgtcagaggtgaaattcttggatttgttaaagacggactactgcgaaagcatttgccaaggatgttttca
>6c9c00178e26af4f55cdc5d799c60924a36a8d34_347
agctccaatagcgtatattaaagttgttgcggttaaaaagctcgtagttggatttctgctaaggacgaccggtccgccttcgggtgtgtacatgtgttgaccgaggcatcaatctggaagacgcctgtacttaaccgtgcgggcgtaattcagacgttttactttgaggaaattagagtgtttcaagcaggcaattgccctgaatacattagcatggaattataaaataggacttcggttctattttgttggtttctagaactgaagtaatgattaatagggacagtcgggggcattcgtactcaactgtcagaggtgaaattcttggatttgttgaagacgaactactgcgaaagcatttgccaaggatgttttca
>201eb31bc4b3a1b0baa7e7292f806802eac14086_260
agctccaatagcgtatattaaagttgttgcggttaaaacgctcgtagtctgcttacagcactggctgccgcttctcagcggacgagtcatgctatggaagcttcttcatctgatgaagcttacattactttgaggaaattagggtgtttaaagcaagcgaacgcttggatattttagcatgggataataacgtaggttctctctcctacttttttggtatgtcggagtgaggaatgtgcaatagggacagttgggggcattcgtacttaacagtcagaggtgaaattcttggatttgttaaagacgaacaactgcgaaagcatctgccaaggatgttttct
>d105b1a03ea109b6b1ccc24748fa4a07f792f9c3_322
agctccaatagcgtatattaaagttgttgcagttaaaaagctcgtagttgaatttctgctggagttgagtaggtctgctctttgagtgtgtaccttgtttcgacttcgcatcctcttagaatgttattaaagtactttattgtgctttatataatatctaagatttttactttgagtaaattagagtgtttcaagcaggcatattgccttgaatactttagcatggaataataagataggattttggttctattttattggtttatagaactaaagtaatgattaatagggacagttgggggcattcgaatttaactgtcagaggtgaaattcttagatttgttaaagtcgaactactgcgaaagcatttgccaaggatgttttca
>a8d17c65edb97723b4ebe56373c07ee0a8b1567f_354
agctccaatagcgtatattaaagttgttgcggttaaaaagctcgtagttggatttctgccgaggacgaccggtccgccctccgggtgtgcatctggatcggcctgggcatcttcttggagaacgtatctgcacttcattgtgtggtgcggtatccaggacctttactttgaggaaattagagtgtttcaagcaggcacacgccttgaatacattagcatggaataataagataggaccttggttctattttgttggtttctagaactgaggtaatgattaatagggatagttgggggcattcgtatttaactgtcagaggtgaaattcttggatttgttaaagacggactactgcgaaagcatttgccaaggatgttttca
>a016c95a288d05ea8b27c2acb3dcbe4764f2e709_229
agctccaatagcgtatattaaagttgttgcggttaaaaagctcgtagttggatttctgccgaggacgaccggtcggccctctgggtttgtatctggctcggcctgggcatcttcttggagaacgtaaccgcacttgactgtgtggtgcggtattcaggacttttactttgaggaaattagagtgtttcaagcaggcacacgccttgaatacattagcatggaataataagataggacctcggttctattctgttggcttctagagctgaggtaatgattgatagggatagttgggggcattcgtatttaactgtcagaggtgaaattcttggatttgttaaagacggactactgcgaaagcatttgccaaggatgttttca
>99753a43a096b4ef9366c5e533d583784e84c3ad_158
agctccaatagcgtatattaaagttgttgcggataaaaagctcgtagttggatttttgccgagaacgactggtccgccctctgggtgagtatctagctcggtctgggcatcttcttggagaacgtagctgcacttgactgtgtggtgcggtatccaggacttttactttgaggaaattagagtgtttcaagcaggcgcacgccctgaatacattagcatggaataataagataggaccttggttctattttgttggtttctagagctgaggtaatgattaatagggatagttgggggcattcgtatttaactgtcagaggtgaaattcttggatttgttaaagacggactactgcgaaagcatttgccaaggatgttttca
>2c737f66a2f1b0de54dfda8fa50bd7f42e874358_180
agctccaatagcgtatattaaagttgttgcagttaaaaagctcgtagttgaatttctgctggagttgaataggtctgctcgatgagtgtgtaccttgttttcgacttcgcatcctcttagaatgttattaaagtactttattgtgctttaaataatatctaagatttttactttgagtaaattagagtgtttcaagcaggcatattgccttgaatactttagcatggaataataaaataggattttggttctattttattggtttatagaactaaagtaatgattaatagggacagttgggggcattcgaatttaactgtcagaggtgaaattcttagatttgttaaagtcgaactactgcgaaagcatttgccaaggatgttttca
>739bd864eecb94c6de0a292846ccbaf18f4240a7_170
agctccaatagcgtatattaaagttgttgcggttaaaaagctcgtagttggatttctgtcgaggacgatcggtccgccttctgggtgagtatctggctcggccttggcatcttcttggagaacgtactgcacttcactgtgtggtgcggaatccaggacctttactttgaggaaattagagtgtttcaagcaggcgcacgccttgaatacattagcatggaataataagataggacctcggttctattttgttggtttctagagctgaggtaatgattaatagggatagttgggggcattcgtatttaactgtcagaggtgaaattcttggatttgttaaagacggactactgcgaaagcatttgccaaggatgttttca
>5e37693be6eb6f802cd812fca22107248e61e6c7_302
agctccaatagcgtatattaaagttgttgcggttaaaaagctcgtagttggatttttcatagttcgtatacttcagtatagctgaattatgatattaaagattttcgttaatttgttactttgaggaaattagagtgtttaaagcagatttatgtcttgtatatattagcatggaataacattaaggattaaaagaactttgttggttactttttttaataatgtttgatagggacaaccgggggcatttgtatttaacagtcagaggtggaattcttggatttgttaaggacaaacaattgcgaaagcatttgccaaggatgttttcg
>43fa085caa566a36e25969e2e97ffdea31eda045_129
agctccaatagcgaatattaaagttgttgcggttaaaaagctcgtagttggatttctgcttaggacgaccggtccgcccactgggtgagtatctggctcggcctgggcatcttcttggagaacgtagctgcgcttgattgtgtggtgcggaatccaggacttttactttgagaaaattagagtgtttcaagcaggcacacgccttgaatacattagcatggagtaataatataggacctcggttctattttgttggtttctagagctgaggtaatgattgatagggatagttggtggcattcgtatttaactgtcagaggtgaaattcttggatttgttaaagacggactactgcgaaagcatttgccaaggatgttttca
>01a8638e3f30ab7846a50539a306dc52fdf82dbd_119
agctccaatagcgtatattaaagttgttgcggttaaaaagctcgtagttggatttctgccaaggatgaccggtccgccttcgggtgtgtacctgtgttgtccgaggcatctttctggttggacgccagcacttcaatgtgttgtcgtactccagaacttttactttgaggaaattagagtgtttcaagcaggcatttgccttgaatacattagcatggaataataatctaggactccggttctattttgttggttctgagaacaggagtaatgattaatagggacagtcgggggcattcgtactcaactgtcagaggtgaaattcttggattagttggagacgaactactgcgaaagcatttgccaaggatgttttca
>a06026dd39a4ca9d4009ad2d758ae0aaf0cdc792_147
agctccaaaagcgtatattaaagttgttgcggttaaaaagctcgtagttggattttatagtcgaagctacattttggtgtaactcgattatggtaataatttattattacacgttactttgaggaaattagagtgtttaaagcagatctttgtcatgtatatattagcatggaataacactaaggattattaaaactttgttggttactttttttaataatgattgatagggacagccgggggcatttgtatttaacagtcaggaattcttggatttgttaaggacaaactattgcgaaagcatttgccaaggatgttttcg
>2da5b3eb55017c94264e866e07818b0864a5c3c2_176
agctccaatagcatatattaaagttgttgcggttaaaaagctcgtagttggatttctgctgcagatgaccggtccgccttccgggtgagtatctggctcgtctgtggcattttcttggagaacgtgtctgcacttgactgtgtggtgcgccatctgggacttttactttgaggaacttagagtgcttcaagcaggccagtgccttgaatacattagcatggaataataagataggaccttggttttattttgctggtttctaaaactgaggtaaggattgatagggacagttgggggcattcgtacttaactgtcagaggtgaaattcttggatttgttaaagacggaccactgcgaaagcatttgccaaagatgttttca
>e04dc0f62a64ef583cc4dc4c706a680b2e1aa6eb_162
agctccaatagcgtatattaaagttgttgcggttaaaaaactcgtagttggatttctgccgaggacgaccggtccgccctccgggtgtgcatctggatcggcctgggcatcttcttggagaacgtatctgcacttcattgtgtggtgcggtatccaggacctttactttgaggaaattagagtgtttcaagcaggcacacgccttgaatacattagcatggaataataagataggaccttggttctattttgttggtttctagaactgaggtaatgattaatagggataggtgggggcattcgtatttaactgtcagagttgaaattcttggatttgttaaagacggactactgcgaaagcatttgccaaggatgttttca
>829d1b164c2896b8c6060a79bb6895fb586bf64f_129
agctccaatagcgtatattaaagttgttgcggttaaaaagctcgtagtcgctcttctgtcagaaacgaccggtccgccctttgggtgagtatctggctcggctctagcctcttcttggttgaccatgctgcacttgagtgtgtggcaagggacccaggactttcactttgagaaaattggagtgtttcaagcaggctcacgccgtggacacataagcatggaatgaatggtttgaacctcgcgtctatcttgttggtttctagaaatggggtgatcaagatagggatagttgggggtattcgtatttaactgtcagaggtgaaattcttggatttgttaaagacggaccactgcgaaagcatttgccaaagatgttctca
>844962070437fe7f89b7dc2c17e08eb06d7ed1c7_233
agctccaatagcgtatattaaagttgttgcggttaaaaagctcgtagttggatttctgctgaggacgatcggtccgccctctgggtgagtatctggctcggccttggcatcttcttggagaacgtagctgcacttgactgtgtggcgcggtatccaggacttttactttgaggaaattagagtgtttcaagcaggcgcacgccttgaatacattagcatggaataataagataggacctcggttctattttgttggtttctagagctgaggtaatgattaatagggatagttgggggcattcgtatttaattgtcagaggtgaaattcttggatttgttaaagacggactactgcgaaagcatttgccaaggatgttttca
>45f8434a67b7601bc476b51e5d4be9b668e6a919_135
agctccaatagcgtatattaaagttgttgcagttaaaaagctcgtagttggatttctgtccaaaatgaccggtccaccctcgtggtgagacagggttcgatttggacattttcctgaggaacgcgctgcacttcactgtgtggtcggaattcaggacttttactttgagaaaattagagtgtttcaagcaggcctacgctttgaatactgcagcatggaataatatgataagacttcggttttattttgtgggtttctaaaactgaagtaatgattaatagggacagttgggggcattcgtatttaactgtcagaggtgaaattcttggatttgttaaagacgaactactgcgaaagcatttgccaaggatgttttca
>f90e1d3417861d7ea72b0374db53e0232b7c069e_116
agctccaatagcgtatattaaagttgttgcggttaaaaagctcgtagttggatttctgctgaggacgatcggtccgccctctgggtgagtatctggctcggccttggcatcttcttggagaacgtagctgcacttgactgtgtggcgcggtatccaggacttttactttgaggaaattagagtgtttcaagcaagcgcacgccttgaatacattagcatggaataataagataggacctcggttttattttgttggtttctagagctgaggtaatgattgatagggatagttgggggcattcgtatttaactgtcagaggtgaaattcttggatttgttaaagacggactactgcgaaagcatttgccaaggatgttttca
>0977faa48e661e7fcc5cef43d3ffa6b9d25c2452_139
agctccaatagcgtatattaaagttgttgcggttaaaaagctcgtagttggatttctgccaaggacgaccggtccgccttcgggtgtgtatctgtgttgaccgaggcatctttctaggtgacgcccgtgcttcactgcacggagcgtgctctagaacttttactttgaggaaattagagtgttccaagcaggcctatgccctgaatacattagcatggaataataaggtaggacttcggttctattttgttggtttctagaactgaagtaatgattaatagggacagtcgggggcattcgtactcaactgtcagaggtgaaattcttggatttgttggagacgaactactgcgaaagcatttgccaaggatgttttca
>5b69eb05cc2d757a032e3e8e78d0246e529ed185_198
agctccaatagcgtatattaaagttgttgcggttaaaaagctcgtagttggatttctgccgaggacgaccggtccgccctctgggtgagcatctggttcggcctgggcatcttcttggagaacgtcgctgcacttgactgtgtggtgcggtatccaggacttttactttgaggaaattagagtgtttcaagcaggcacacgccttgaatacattagcatggaataataagataggacctcggttctattttgttggtttctagggctgaggtaatgattaatagggatagttgggggcattcgtatttaactgtcagaggtgaaattcttggatttgttaaagacggactactgcgaaagcatttgccaaggatgttttca
>8b4c6ec50c3741216da330b2ebba6694177e2658_96
agctccaatagcgtatattaaagttgttgcagttaaaaagctcgtagttgaatttctgcagaatgcgacgcgatgcaccttctaggcgatttcgtgtttgcttttgcattctggtgactgctgtatacacttaactgtttatactgctgttcactagcatttactttgagaaaattagagtgtttcaagcaggctttttgccttgaatttaactgtcagaggtgaaattcttagatttgttaaagacgaaccaatgcgaaagcatttgccaaaaatgttttca
>1695c4a0bf3dbde87ef638bf0b6f4da715b657f7_105
agctccaatagcgtatattaaagttgttgcggttaaaaagctcgtagttggatttctgccgaggacgaccggtccgccctccgggtgtgcatctggatcggcctgggcatcttcttggagaacgtatctgcacttcattgtgtggtgcggtatccaggacctttactttgaggaaattagagtgtttcaagcaggcacacgccttgaatacattagcatggaataataagataggaccgtggttctattttgttggtttctagaactgaggtaatgattaataggaatagttgggcgcattcgtatttaactgtcagaggtgaaattcttggatttgttaaagacggactactgcgaaagcatttgccaaggatgttttca
>c523a186a8e841fb4c4a521e3dbd20bd14c86c4d_107
agctccaatagcgtatattaaagttgttgcggttaaaaagctcgtagttggatttctgccgaggacgaccggtccgccctccgggtgtgcatctggatcggcctgggcatcttcttggagaacgtatctgcacttcattgtgtggtgcggtatccaggacctttactttgaggaaattaaagtgtttcaagcaggcacacgccttgaatacattagcatggaataataagataggaccttggttctattttgttggtttctagaactgaggtaatgattaataggaatagttgggggcattcgtatttaactgtcagaggtgaaattcttggatttgttaaagacggactactgcgaaagcatttgccaaggatgttttca
>8b58a06f94ea2c4d9c3465d64311ad760dde1594_176
agctccaatagcgtatattaaagttgttgcggttaaaaagctcgtagttggatttaacaattcaatttttacttcggtaattaattgttttggttttagtgactttgtcactaaattgttactttgaggaaattagagtgtttaaagcagatttatgtcatgaatatattagcatggaataacattaaggattaagagaactttgttggttacttcttttaataatgtttgatagggacagccgggggcatttgtatttaacagtcagaggtggaattcttggatttgttaaggacaaactattgcgaaagcatttgccaaggatgttttcg
>e823fb81d263ce8786e75badb0f0f44bc52e5e6e_78
agctccaatagcgtatattaaagttgttgcggttaaaaagctcgtagttggatttctgcttagggtgaccggtccgccttcgggtgtgtatcagtgttgtccaaagcatctgtctagttgcgctcgtgcttcaatgcacgctgcgtgttctagaccttttactttgaggaaattagagtgtttcaagcaggcatatgccttgaatacattagcatggaataataatttaggacttcggttctattttgttggtttctagaactgaagtaatgattaatagggacaatcgggggcattcgtactcaacagtcagaggtgaaattcttggatttgttggagacgaactactgcgaaagcatttgccaaggatgttttca
>1bd194b0d0e51088ec189bae6487b604e14c0ba0_105
agctccaatagcgtatattaaagttgttgcggttaaaaagctcgtagttggatttctgctgaggacggccggtctgcccaccgggtatgcatctggctcggcctgagcatcttcctggcgaacgccgctgcacttcattgtgtggcacggtatccaggacttttactttgaggaaattagagtgcttaaagcaggcatacgcctgaacacatgagcatggaataatgtgataggacctcggttctattttgttggtttctagagctgaggtaatgataaatagggacagttgggggcattcgtatttaactgtcagaggtgaaattcttggatttgttaaagacggaccactgcgaaagcatttgccacggatgtcttca
>3a7d4ab1008d9a1bb28838c522ade02ff3ec0374_83
agctccaatagcgtatattaaagttgttgcagttaaaaagctcgtagttgaatttctgcgggagttgaataggtctgctctttgagtgtgtaccttgtttcgattccgcatcctcttagaatgttattaaagtactttattgtgctttatataatatctaagatctttactttgagtaaattagagtgtttcaagcaggcatattgccttgaatactttagcatggaataataagataggattttggttctattttattggtttatagaactaaagtaatgattaatagggacagttgggggcattcgaatttaactgtcagaggtgaaattcttagatttgttaaagtcgaactactgcgaaagcatttgccaaggatgttttca
>a8aca37ba2936ac92ae64b658a3b595eff025d36_88
agctccaatagcgtatattaaagttgttgcggttaaaaagctcgtagttagatttctgatttagtcgaccggtcgttcctttaatggaatgtatcaggtttgattaaattatttgtctgattatttctggctgcacttaactgtgtggctggccattcagaccatttactttgaggaaattagagtgtttcaagcaggcttacgccttgaatacgttagcatggaataataaaataagactttggttttattttgttggtttctaaaactaaagtaatgattaatagggatagtcgggggcattcgtacttaactgtcagaggtgaaattcttagatttgttaatgacgaactactgcgaaagcatttgccaaggatgttttca
>756c4c1a7932c3553c365ddb070b125c1c1235d5_82
agctccaatagcgtatattaaagttgttgcggttaaaaagctcgtagttggatttctgttgaggacgatcggtccgccttctgggtgagtatctggctcggccttggcatcttcttggagaacgtagtgcacttcactgtgtggtgcggaatccaggacctttactttgaggaaattagagtgtttcaagcaggcgcacgccttgaatacattagcatggaataataagataggacctcggttctattttgttggtttctagagctgaggtaatgattaatagggatagttgggggcattcgtatttaactgtcagaggtgaaattcttggatttgttaaagacggactactgcgaaagcatttgccaaggatgttttca
>df7f3711309eca696c2529a44b78f4e6de0badb3_67
agctccaaaagcgtatattaaagttgttgcggttaaaaagctcgtagttggattttatagtcgaagcaacattttggtgtaactcgattatggtaataatttattattacacgttactttgaggaaattagagtgtttaaagcagatctttgtcatgtatatattagcatggaataacactaaggattattaaaactttgttggttactttttttaataatgattgatagggacagccgggggcatttgtatttaacagtcagagatggaattcttggatttgttaaggacaaactattgcgaaagcatttgccaaggatgttttcg
>434335cd1d366058a4e591b3ff7902ba3d712f90_62
agctccaatagcgtatattaaagttgttgcggttaaaaagctcgtagttggatttctgctgaggcctaacggtcagcccattgggtttgcatctgctacggcctgagcatcttcttggagaaggttgctgcacttgactgtgtggcgcggtatccaggacttttactttgaggaaattagagtgcttcaagcaggcacacgccttgaacacgtgagcatggaataatgtgataggacctcggttctattttgttggtttctagagctgaggtaatgataaatagggacagttgggggcattcatatttaactgtcagaggtgaaattcttggatttgttaaagatggactactgcgaaagcatttgccacggatgtcttca
>6e0e2138c79115b5b58f2a77f927f174228d7a17_69
agctccaatagcgtatattaaagttgttgcggttaaaaagctcgtagttggatttctgccgaggacgaccggtccgcccttgcggtgagtatctggctcggcctgggcatcttcttggagaacgccaccacacttgactgtgtggtgcggtatccaggacttttactttgaggaaattagagtgtttcaagcaggcacacgccttgaatacattagcatggaataatatgataggacctcggttctattctgttggcttctagagctgaggtaatgattgatagggatagttgggggcattcgtatttaactgtcagaggtgaaattcttggatttgttaaagacggactactgcgaaagcatttgccaaggatgttttca
>b2d640bf8622b47281b523242ee8a660e8615ec5_58
agctccaaaagcgtatattaaagttgttacggttaaaaagctcgtagttggattttatagtcgaagctacattttggtgtaactcgattatggtaataatttattattacacgttactttgaggaaattagagtgtttaaagcagatctttgtcatgtatatattagaatggaataacactaaggattattaaaactttgttggttactttttttaataatgattgatagggacagccgggggcatttgtatttacagtcagaggtggaattcttggatttgttaaggacaaactattgcgaaagcatttgccaaggatgttttcg
>2172d83cfe1ad4481c6dc8941e3c7a98f6ab27f5_70
agctccaatagcgtatattaaagttgttgcagttaaaaagctcgtagttgaatttctgctggagttgattaggtctgctctatgagtgtgtaccttgtttcgacttcgcatcctcttagaatgttattaaagtactttattgtgctttatataatatctaagatctttactttgagtaaattagagtgtttcaagcaggcatattgccttgaatactttagcatggaataataaaataggactttggttctattttattggtttatagaactaaagtaatgattaatagggacagttgggggcattcgaatttaactgtcagaggtgaaattcttagatttgttaaagtcgaactactgcgaaagcatttgccaaggatgttttca
>f07d92a984474455dd525fd737a728391a2b6bb6_86
agctccaatagcgtatattaaagttgttgcggttaaaaagctcgtagttagatttttgatctagtcgaccagtcactccaatggaatgtatcaggtttgattagattatttgtctgaaatttttgtctgcacttcactgtgtggtcaaagattcagaccttttactttgaggaaattagagtgtttcaagcaggcatttgccttgaatacgttagcatggaataataaaataagactttggttttattttgttggtttctaaaactaaagtaatgattaatagggatagtcgggggcattcgtacttaactgtcagaggtgaaattcttggatttgttaaagacgaactactgcgaaagcatttgccaaggatgttttca
>5840d9030eea063cf24dddc0d0f969885caff52e_62
agctccaatagcgtatattaaagttgttgcggttaaaaagctcgtagttggatttctgccgaggacgaccggtccgccctccgggtgtgcatctggatcggcctgggcatcttcttggagaacgtatctgcacttcattgtgtggtgcggtatccaggacctttactttgaggaaattagagtgtttcaagcaggcacacgccttgaatacattagcatggaataataagataggaccttggttatattttgttgggttctagaactgaggtaatgattaatagggatagttgggggcattcgtatttaactgtcagaggtgaaattcttggatttgttaaagacggactactgcgaaagcatttgccaaggatgttttca
>22fc98e5e89a61cd4562042d0c02fe16ec2359f3_82
agctccaatagcgtatattaaagttgttgcggttaaaaagctcgtagttggatttctgccaaggatgaccggcccgccttcgggtgtgtgcatgcgttgaccaaggcattttcttgttattgatatgactttactgccatatcaatgttcaagaacgttactttgaggaaattagagtgtttcaagcaggcaatcgccctgaatacattagcatggaataacaatataggactctggttctattttgttggtttctagagctggagtaatgattaatagggacaatcgggggcatccgtactcaaccgtcagaggtgaaattcttggattggttgaagacgaactactgcgaaagcatttgccaaggatgttttca
>bd48536b13019496a7df289722c32a7199266219_107
agctccaatagcgtatattaaagttgttgcggttaaaaagctcgtagttggatttctgctaaggatgaccggcccgccttcgggtgtgtgcatgtgttgaccgaggcatttttctgatattgacttcacttaactgggttgtcaattttcagagatgttactttgaggaaattagagtgtttcaagcaggcaattgccctgaatacattagcatggaataacaatataggactccggttctattttgttggtttcttgagctggagtaatgattaatagggacaatcgggggcatccgtactcaaccgtcagaggtgaaattcttggattggttgaagacgaactactgcgaaagcatttgccaaggatgttttca
>941c0fa051a096e003c699f7747c41bf07765314_82
agctccaatagcgtatattaaagttgttgcggttaaaaagctcgtagttggatttctgccgaggacgaccggtccgccctctgggtgcgtatctggctcggcctgggcatcttcttggagaacgtatctgcacttgactgtgtggtgcggtatccaggacttttactttgaggaaattagagtgtttcaagcaggcatacgccttgaatacattagcatggaataataagataggacctcggttctattttgttggtttctagagctgaggtaatgattaatagggatagttgggggcattcgtatttaactgtcagaggtgaaattcttggatttgttaaagacggactactgcgaaagcatttgccaaggatgttttca
>6868d656630344ab420014389943aece04921224_101
agctccaatagcgtatattaaagttgttgcggttaaaaagctcgtagttggatttctgctgaggacgaccggtccgccctctgggtgagtatctggctcggccttggcatcttcttggagaacgttactgcacttgattgtgtggtgcggtatccaggacttttactttgaggaaattagagtgtttcaagcaggcgcacgccttgaatacattagcatggaataataagataggaccttggttctattttgttggtttctagagctgaggtaatgattaatagggatagttgggggcattcgtatttaactgtcagaggtgaaattcttggatttgttaaagacggactactgcgaaagcatttgccaaggatgttttca
>c71b4f6b6fea023459721b3e84a267ce1e4f3d54_55
agctccaatagcgtatattaaagttgttgcggttaaaaagctcgtagttggatttctgcttaggacgaccggtccgcccactgggtgagtatctggctcggcctgggcatcttcttggagaacacaactgcacttgactgtgtggtgtggtatccaggacttttactttgagaaaattagagtgtttcaagcaggcacacgccttgaatacattagcatggaataataatataggacctcggttctattttgttggtttctagagctgaggtaatgattgatagggatagttgggggcattcgtatttaactgtcagaggtgaaattcttggatttgttaaagacggactactgcgaaagcatttgccaaggatgttttca
>62b0f924cadcaac286ef0e17d80c84d052b77911_55
agctccaatagcgtatattaaagttgttgcagttaaaaagctcgtagttgaattgaaggttaaatgtgcttgtgaattaggccaaaaagcttagttcacaggtcatacctgatgtttaggttgatatcctgcacttaattgggtggggtatttcgacttagaccgtttaccttgaagaaattggagtgtttaaagcaggcgtttcgtttgaacatgttagcatgggataatgaaatatgacatctacatattcgttggcgtatacttgtaggtgtaatgatggataggggtagcgggagtggttggtatttagcggctagaggtgaaattcttagattcgctaaagactcacagaagcgaaagcgttccacgacaatacccccg
>8a9d5714f75cc03908f4a56d6294f4d7a565b8ba_77
agctccaatagcgtatattaaagttgttgcggttaaaaagctcgtagttggatttctgccaaggatgaccggcccgccttcgggtgtgtgcatgtgttgaccgaggcatttttctgatgttgacttcacttcactgggttgtcaattttcagagatgttactttgaggaaattagagtgtttcaagcaggcaattgccctgaatacattagcatggaataacaatataggactccggttctattttgttggtttcttgagctggagtaatgattaatagggacaatcgggggcatccgtactcaaccgtcagaggtgaaattcttggattggttgaagacgaactactgcgaaagcatttgccaaggatgttttca
>406d953c7717cabc7d0b33cc312c35583a3f6e92_48
agctccaatagcgtatatttaagttgttgcagttaaaaagctcgtagttggatttctgcttaggacgaccggtccgcccttgagcatctggctcggcctgggcatcttcttggagaacgtagctgcgcttgattgtgtggtgcggaatccaggacttttactttgagaaaattagagtgtttcaagcaggcacacgccttgaatacattagcatggaataataatataggacctcggttctattttgttggtttctagagctgaggtaatgattgatagggatagttgggggcattcgtatttaactgtcagaggtgaaattcttggatttgttaaagacggactactgcgaaagcatttgccaaggatgttttca
>1a33ca03fc0df5c880a38e8955b39770bb2f86a4_54
agctccaatagcgtatattaaagttgttgcggttaaaaagctcgtagttggatttctgctgaggacgaccggtccgccctctgggtgagtatctggctcggccttggcatcttcttggagaacgtaactgcacttgactgtgtggtgcggtatccaggacttttactttgaggaaattagagtgtttcaagcaggcgcacgccttgaatacattagcatggaataatgagataggaccttggttctattttgttggtttctagagctgaggtaatgattaatagggatagttgggggcattcgtatttaactgtcagaggtgaaattcttggatttgttaaagacggactactgcgaaagcatttgccaaggatgttttca
>a512cec226db446585066f21cb2ee78eb7909a02_47
agctccaatagcgtatattaaggttgttgcggttaaaaagctcgtagttggatttctgctgaggacgaccggtccgccctctgggtgagtatctggctcggccttggcatcttcttggagaacgttactgcacttgattgtgtggtgcggtatccaggacttttactttgaaaaaattagagtgtttcaagcaggcgttttgctatgaatacattagcatggaataataagataggacctcggttctattttgttggtttctagagctgaggtaatgattaatagggatagttgggggcattcgtatttaactgtcagaggtgaaattcttggatttgttaaagacggactactgcgaaagcatttgccaaggatgttttca
>25a28d0babb0d60e9540d8854c9d6c8565de2d7b_50
agctccaaaagcgtatattaaagttgttgcggttaaaaagctcgtagttggattttatagtcgaagctacattttggtgtaactcgattatggtaataatttattattacacgttactttgaggaaattagagtgtttaaagcagatctttgtcatgtatatattagcatggaataacactaaggattattaaaactttgttggttacttttttttaataatgattgatagggacagccgggggcatttgtatttaacagtcagaggtggaatttttggatttgttaaggacaaactattgcgaaagcatttgccaaggatgttttcg
>f07fe2f6863eaffd02c34329d2fe3d42a6bda1f0_46
agctccaatagcgtatattaaagttgttgcggttaaaaagctcgtagttggatttctgttgaggacgatcggtccgccttctgggtgagtatctggctcggccttggcatcttcttggagcgcgtactgcacttcattgtgtggtgcgaaatccaggacctttactttgaggaaattagagtgtttcaagcaggcgcacgccttgaatacattagcatggaataataagataggacctcggttctattttgttggtttctagagctgaggtaatgattaatagggatagttgggggcattcgtatttaactgtcagaggtgaaattcttggatttgttaaagacggactactgcgaaagcatttgccaaggatgttttca
>324400ced585bf75f1c2890a951aba1a4d6eb460_58
agctccaatagcgtatattaaagttgttgcagttaaaaagctcgtagttgaatttctgctggagttgaataggtctgctctttgagtgtgtaccttgtttcgacttcgcatcctcttagaatgttattaaagtactttattgtgctttatataatatctaagatttttactttgagtaaattagagtgtttaaagcaggcatattgccttgaatactttagcatggaataataaaataggactttggttttattttattggtttatagaactaaagtaatgattaatagggacagttgggggcattcgaatttaactgtcagaggtgaaattcttagatttgttaaagtcgaactactgcgaaagcatttgccaaggatgttttca
>e398752440eed639b239a26a925e360b69b299f8_191
agctctgatagtatatattaaagttgttgcggttaaaaagctcgtagttggatttctgcttaggacgaccggtccgcccactgggtgagtatctggctcggcctgggcatcttcttggagaacgtagctgcgcttgattgtgtggtgcggaatccaggacttttactttgagaaaattagagtgtttcaagcaggcacacgccttgaatacattagcatggaataataatataggacctcggttctattttgttggtttctagagctgaggtaatgattgatagggatagttgggggcattcgtatttaactgtcagaggtgaaattcttggatttgttaaagacggactactgcgaaagcatttgccaaggatgttttca
>141fb7981938a04cae738acc6e1a4461178c37bf_49
agctccaagagcgtatattaaagttgttgcggttaaaaagctcgtagttggatttctgccgaggacgaccggtccgccctctgggtgagtatctggctctgcctgggcatcttcttggagaacgtagctgcacttcactgtgtggtgcggtatccaggacttttactttgaggaaattagagtgtttcaagcaggcatacgccttgaatacattagcatggaataataagataggacctcggttctattttgttggtttctagagctgaggtaatgattaatagggatagttgggggcattcgtatttaactgtcagaggtgaaattcttggatttgttaaagacggactactgcgaaagcatttgccaaggatgttttca
>717ab87a0c314e50ef5a691d5834310a412134fe_47
agctccaatagcgtatattaaagttgttgcggttaaaaagctcgtagttggatttctgcttaggacgaccggtccgcccactgggtgagtatctggctcggcctgggcatcttcttggagaacgtagctgcgcttgattgtgtggtgcggaatccaggacttttactttgagaaaattagagtgtttcaagcaggcacacgccttgaatacattagcatggaataatgagatagggccttgatggaagacgtcatgtctattttgttggtttgcacgccaaggcaatgattgacagggatagttgggggcattcgtatttaactgtcagaggtgaaattcttggatttgttaaagacggactactgcgaaagcatttgccaaggatgttttca
>04acab29ba78ca3d301602c7083326ca9fb8268d_35
agctccaatagcgtatattaaagttgttgcggttaaaaagctcgtagttggatttctgcttaggacgaccggtccgcccactgggtgagtatctggctcggcctgggcatcttcttggagaacgtagctgcgcttgattgtgtggggcggaatccaggacttttactttgagaaaattagagtgtttcaagcaggcacacgccttgaatacattagcatggaataataatataggacctcggttctattttgttggtttctagagctgaggtaatgattgatagggatagttgggggcattcgtatttaactgtcagaggtgaaattcttggatttgttaaagacggacaaatgcgaaagcatttgccaaggatgttttca
>557eccbbd5a730515faee95c3fd93332d424bb1a_43
agctccaatagcgtatattaaagttgttgcggttaaaaagctcgtagttggatttctgccgaggacgatcggtccgccctctgggtgcgtatctggctcggcctgggcatcttcttggagaacgtatctgcacttgactgtgtggtgcggtatccaggacttttactttgaggaaattagagtgtttcaagcaggcatacgtcttgaatacattagcatggaataataagataggacctcggttctattttgttggtttctagagctgaggtaatgattaatagggatagttgggggcattcgtatttaactgtcagaggtgaaattcttggatttgttaaagacggactactgcgaaagcatttgccaaggatgttttca
>b50db50ae4943641b8a86e73a9dc95baa4a19980_79
agctccaatagcgtatattaaagttgttgcggttaaaaagctcgtagttggatttctgcttaggatgaccggtccgccttcgggtgtgtatcagcgttgtccagagcatctttctagttgcgctcgtgcttcaatgtacgatgcgtgttctagatcttttactttgaggaaattagagtgtttcaagcaggcatatgccttgaatactttagcatggaataataagataggacttcggttctattttgttggtttctagaactgaagtaatgattaatagggacaatcgggggcattcgtactcaacagtcagaggtgaaattcttggatttgttggagacgaactactgcgaaagcatttgccaaggatgttttca
>bea5d30af60fb8467c4e1d6ec2491c7a027f34ae_46
agctccaatagcgtatattaaagttgttgcggttaaaaagctcgtagttggatttctgctgaggacgaccggtccgccctctgggtgagcatctggttcggcctgagcatcctcttggagaaggcttgcgcacttcactgtgtgcgaccgtatccaggacttttactttgaggaaattggagtgcttcaagcaggcatacgccttgaacacatgagcatggaataatgcgataggacctcggttctattttgttggtttctagagctgaggtaatgataaatagggacagttgggggcattcgtatttaactgtcagaggtgaaattcttggatttgttaaagacggaccactgcgaaagcatttgccacggatgtcttca
>de7add9cb1d2ef7f2b25e550b24a9fdee05fefb0_41
agctccaatagcgtatattaaagttgttgcggttaaaaagctcgtagttggatttctgccaaggatgaccagtccgccttcgggtgtgtatctgtgttgaccgcggcatcttcctaggaaacgtctgtacttaactgtacggcacgtattctaggacttttactttgaggaaattagagtgtttcaagcaggcatatgccttgaatacattagcatggaataatgaggtaggactccggttctattttgttggtttctagaacaggagtaatgattaatagggacagtcgggggcattcgtactcaactgtcagaggtgaaattcttggatttgttggagacgaactactgcgaaagcatttgccaaggatgttttca
>01f843ca567112849d23759e6fc29b0c6c8b7940_42
agctccaatagcgtatattaaagttgttgcggttaaaaagctcgtagttggacttctgcgaaggacgaccggcccgcccaccgggtgagtatctggttcggcctttgcatcctcttggagaaggtgtgttcacttcactgtgtgcacccgtatccaggacattttactttgaggaaattggagtgcttcaggcaggcacacgccttgaacacatgagcatggaataatgcgataggacctcggttctattttgttggtttctagcactgaggtaatgataaatagggacagttgggggcattcgtatttaactgtcagaggtgaaattcttggatttgttaaagacggaccactgcgaaagcatttgccacggatgtcttca
>de73d3a95a35d48bee80d3e42f5f428675979f9c_37
agctccaatagcgtatattaaagttgttgcggttaaaaagctcgtagttggatttctgctgaggacgaccggtccgccctctgggtgagtatctggctcggccttggcatcttcttggagaacgttactgcacttgattgtgtggtgcggtatccaggacttttactttgcggaaattagagtgtttcaagcaggcgcacgccttgaatacattagcatggaataataagataggacctcggttctattttgttggtttctagtactgaagtaatgattgatagggatagttgggggcattcgtatttaactgtcagaggtgaaattcttggatttgttaaagacggactactgcgaaagcatttgccaaggatgttttca
>1d0d47eb3fed4d61b55a7038a12b1a450c99d492_68
agctccaatagcgtatattaaagttgttgcggttaaaaagctcgtagttggatttctgctgaggatgaccggtccgccctctgggtgagtatctggctcagccttggcatcttcctgaagaacgttgctgcacttgactgtgtggtgcggtatttaggacatttactttgaggaaattagagtgtttcaagcaagcgcacgccttgaatacattagcatggaataataagataggacctcggttctattttgttggtttctagagctgaggtaatgattgatagggatagttgggggcattcgtatttaactgtcagaggtgaaattcttggatttgttaaagacggactactgcgaaagcatttgccaaggatgttttca
>ef2d558f3c53c0960a9e2eb5f0ab267ecdaed011_53
agctccaatagcgtatattagagttgttgcagttaaaacgctcgtagtctcaactgggcactggagctgcgggtccgcccgctcgggcggtactcgcaggtccagggccggtttggcgggcgagtggatggccccgtgccacccacccccccgccgctgttactgtgagaaaatcagagtgctcaaagcaggcgcacgcctagacgcattagcatggaataacacgacacgaccggcgggtctgttttgttggtttccggagccgcggtaatgatgaacaggagcagttgggggcattcgtatttaattgtcagaggtgaaattcttggatttattaaagacgaacggaagcgaaagcgtctgccaaggatgctttca
>57659cfc0f021ef7ddf42433f106c987277d2de8_38
agctccaatagcgtatattaaagttgttgcggttaaaaagctcgtagttggatttctgctgaaagtgaccggtccgccttctgggtgagcatctggtgagcttttggcatccttgtggagaacgtatctgcacttgattgtgtggcacggtatccacgacttttactttgaggaacttagagtgtttcaagcaggcctgcgccttgaatacattagcatggaataataagatgggaccttggttcttttttgttggtttctagagctgaggtaatgattgatagggatagttgggggcattcgtatttaactgtcagaggtgaaattcttggatttgttaaagacggactactgcgaaagcatttgccaaggatgttttca
>79aa111ef40cf85c6927695a2ecbabfc6aacaaf8_40
agctccaatagcgtatattaaagttgttgcagttaaaaagctcgtagttggatttctgttcaggatgaccggcccaacctttggttgtgtgtttggttcagcctgaacattttctcggggaacgtcactgcacttgattgtgtggtacggtatccgagaccgttactttgagaaaattagagtgtttcaagcaagcctacgctttgaatactgcagcatggaataacatgataagacttcggttttattttgtgggtttctaaaactgaggtaatgattaatagggacagttgggggcattcgtatttaactgtcagaggtgaaattcttggatttgttaaagacgaactactgcgaaagcatttgccaaggatgttttca
>a0e21fe41beed01ba2ae497754705b95ffdd06e1_30
agctccaatagcgtatattaaagttgttgcggttaaaaagctcgtagttggatttctgccgaggacgaccggtccgccctctgggtgagtatctggctcggcttgggcatcttcttggagaacgtatctgcacttgactgtgtggtgcggtatccaggacttttactttgaggaaattagagtgtttcaagcaggcatacgccttgaatacattagcatggaataataagataggacctcggttctattttgttggtttctagagctgaggtaatgattaatagggatagttgggggcattcgtatttaactgtcagaggtgaaattcttggatttgttaaagacggactactgcgaaagcatttgccaaggatgttttca
>31e59f4d50eaaa10d406bd23e365453ecda4af54_29
agctccaatagcgtatattaaagttgttgcggttaaaaagctcgtagttggatttctgctgaggacgatcggtccgccctctgggtgagtatctggctcggccttggcatcttcttggagaacgtagctgcacttgactgtgtggcgcggtatccaggacttttactttgaggaaattagagtgtttcaagcaggcatacgccttgaatacattagcatggaataataagataggaccttggttctattttgttggtttctagagctgaggtaatgattaatagggatagttgggggcattcgtatttaactgtcagaggtgaaattcttggatttgttaaagacggactactgcgaaagcatttgccaaggatgttttca
>3d618eac050328b7ab982248e773a35b0833c423_27
agctccaatagcgtatattaaagttgttgcagttaaaaagctcgtagttggatttctgcttaggacgaccggtccgcccactgggtgagtatctggctcggcctgggcatcttcttggagaacgtagctgcgcttgattgtgtggtgcggaatccaggacttttcctttgagaaaattagagtgtttcaagcaggcacacgccttgaatacattagcatggagtaataatataggacctcggttctattttgttggtttctagagctgaggtaatgattgatagggatagttgggggcattcgtatttaactgtcagaggtgaaattcttggatttgttaaagacggactactgcgaaagcatttgccaaggatgttttca
>4c47950e31746d3b14dfdc5b1a8c5812549014dd_39
agctccaatagcgtatattaaagttgttgcagttaaaaagctcgtagttggatttctgccgaggacgaccggtccgcccactgggtgtgtatctggctcggcctgggcatcttcttggagaacgtagctgcacttgactgtgtggtgcggtatccaggacttttactttgaggaaattagagtgtttcaagcaggcacacgccttgaatacattagcatggaataataagataggacttcggttctattttgttggtttctagagctgaggtaatgattaatagggatagttgggggcattcgtatttaactgtcagaggtgaaattcttggatttgttaaagacggactactgcgaaagcatttgccaaggatgttttca
>e24f8a79523089a5e1478ab6d45ae30e95ae92e9_29
agctccaatagcgtatattaaagttgttgcggttaaaaagctcgtagttggatttctgctgaggacaaccggtccgcctactgggtgagtatctggtttggcctctgcattttcttggagaacacaactgcacttgactgtgtggtgtggtatccaggacttttactttgagaaaattagagtgtttcaagcaggcacacgccttgaatacattagcatggaataatactataggactttggttctattttgttggtttctagagctgaggtaatgattgatagggatagttgggggcattcgtatttaaccgtcagaggtgaaattcttggatttgttaaagacggactactgcgaaagcatttgccaaggatgttttca
>131f000d2bac2ca8711b7182375283f33b340896_31
agctctaatagcgtatattaaaattgttgcagttaaaaagctcgtagttgaatttctgtggggtagaataggtctgctctttgagtgtgtacctgatttcgaccctacatcctctcagaatgttattggtgtactttattgtgcatcatttaatatctgagacttttactttgagtaaattagcgtgtttcaagcaggcatttcgccctgaatactttagcatggaataataatttaggactttggttctagtttattggtttatgaactaaagtaatgattaatagggatagttggggacactcgaactcaattgtcagaggtgaaattcttagatttattgaagtcgaactactgcgaaagcatttgtcaaggatgttttca
>94d7e5203b4b5cd437c9ec87770a366c06dcfb15_42
agctccaatagcgtatattaaagttgttgcggttaaaaagctcgtagttggatttctgttgaggacgaccggtccgccctctgggtgagtatctggctcggccttggcatcttcttggggaacgttactgcacttgactgtgtggtgcggtatccaggacttttactttgaggaaattagagtgtttcaagcaggcgcacgccttgaatacattagcatggaataataagataggaccttggttctattttgttggtttctagaactgaggtaatgattaatagggatagttgggggcattcgtatttaactgtcagaggtgaaattcttggatttgttaaagacggactactgcgaaagcatttgccaaggatgttttca
>e8bbe0dbf9a15b5ea9c741c350538a8a84b78834_27
agctccaatagcgtatatttaagttgttgcagttaaaaagctcgtagttggatttctgccgaggacgaccggtccgccctctgggtgagtatctggctcggcctgggcatcttcttggagaacgtatctgcacttgactgtgtggtgcggtattcaggacttttactttgaggaaattagagtgtttcaagcaggcatacgccttgaatacattagcatggaataataagataggacctcggttctattttgttggtttctagatctgaggtaatggttaatagggatagttgggggcattcgtatttaactgtcagaggtgaaattcttggatttgttaaagacggactactgcgaaagcatttgccaaggatgttttca
>04afe92529dea4b166caad7b7a7ace2c4edb2801_33
agctccaggagcgtatattaaagttgttgcagttaaaaagctcgtagttgaatttctgctggagttgaataggtctgctctttgagtgtgtaccttgtttcgacttcgcatcctcttagaatgttattaaagtactttattgtgctttatataatatctaagatctttactttgagaaaattagagtgtttcaagcaggcatattgccttgaatactttagcatggaataataaaataggattttggttctattttattggtttatagaactaaagtaatgattaatagggacagttgggggcattcgaatttaactgtcagaggtgaaattcttagatttgttaaagtcgaactactgcgaaagcatttgccaaggatgttttca
>2d07ccc4f13dfd948ef178c1b9efde6cc8c5b3fa_22
agctccaatagcgtatattaaagttgttgcggttaaaaagctcgtagttggatttctgctgaaagtgaccggtccgccttctgggtgagcatctggtgagcttttggcatccttgtggagaacgtatctgcacttgattgtgtggcacggtatccacgacttttactttgaggaacttagagtgtttcaagcaggcctgcgccttgaatacattagcatggaataataagatgggaccttggttcttttttgttggtttctagaactgaggtaatgatgaatagggatagttgggggcattcgtatttaactgtcagaggtgaaattcttggatttgttaaagacggactactgcgaaagcatttgccaaggatgttttca
>50d0c55a83c434ad490f295f387e09dfc6715e0b_31
agctccaatagcgtatattaaagttgttgcggttaaaaagctcgtagttggacttctgctgaggacgaccggtctgcccactgggtatgcatctggctcggcctgagcatcttcttggcgaacgctgctgcacttcgctgtgtggtgcggtatccaggacttttactttgaggaaattagagtgcttaaagcaggcatacgcctgaacacatgagcatggaataatgtgataggacctcggttctattttgttggtttctagagctgaggtaatgataaatagggacagttgggggcattcgtatttaactgtcagaggtgaaattcttggatttgttaaagacggaccactgcgaaagcatttgccacggatgtcttca
>75ff38e18e6f31328fe328f6f886a0825f11a052_22
agctccaatagcgtatattaaagttgttgcggttaaaaagctcgtagttggatttctgccgaggacgaccggtccgccctctgggtgagtatctggctcggcctgggcatcttcttggagaacgtagctgcacttgactgtgtggtgcggtatccaggacatttactttgaggaaattagagtgtttcaagcaggcatacgccttgaatacattagcatggaataataagataggacctcggttctattttgttggtttctagagctgaggtaatgattaatagggatagttgggggcattcgtatttagctgtcagaggtgaaattcttggatttgttaaagacggactactgcgaaagcatttgccaaggatgttttca
>15801d3a29f7961f425dada58e917cfddcc2ddd6_31
agctccaatagcgtatattaaagttgttgcagttaaaaagctcgtagttggatttctgcttaggacgaccggtccgcccactgggtgagtatctggctcagccttggcatcttcttggagaacgtagctgcgcttgattgtgtggtgcggaatccaggacttttactttgagaaaattagagtgtttcaagcaggcacacgccttgaatacattagcatggaataataatataggacctcggttctattttgttggtttctagagctgaggtaatgattgatagggatagttgggggcattcgtatttaactgtcatagttgaaattcttggatttgttaaagacggactactgcgaaagcatttgccaaggatgttttca
>20953b6e64482764034c2355a3eeac795802a175_30
agctccaatagcgtatattaaagttgttgcagttaaaaagctcgtagttggatttctgttcaggtagaccggtgcactttcgggtgtttacctggtttcgcctggacattttctcggagaacgtcactgcacttcattgggtggtgcggaatccaggacctttactttgagaaaattagagtgtttcaagcaggccaacgctttgaatactgcagcatggaataatatgataagacttcggttttattttgtgggtttctaaaactgaagtaatgatcaatagggacagttgggggcattcgtatttaactgtcagaggtgaaattcttggatttgttaaagacgaactactgcgaaagcatttgccaaggatgttttca
>2a87298f5ccf55068ce47e58101460d7d6635646_23
agctccaatagcgtatattaaagttgttgcggttaaaaagctcgtagttggatttctgctgaggacgaccggtccgccctctgggtgagtatctggctcggccttggcatcttcttggagaacgttactgcacttgattgtgtggtgcggtatccaggacttttactttgaggaaattagagtgtttcaagcaggcgcacgccttgaatacattagcatggaataatagaataggacgtcgtttctattttgttggttttcggaaatcgacgtaatgattaatagggatagttgggggcattcgtatttaattgtcagaggtgaaattcttggatttgttaaagacggactactgcgaaagcatttgccaaggatgttttca
>2e394ea765bf21bafd2b1cb1ffcf23c07e345320_25
agctccaatagcgcatattaaagttgttgcggttaaaaagctcgtagttggatttctgcttaggacgaccggtccgcccactgggtgagtatctggctcggcctgggcatcttcttggagaacgtagctgcgcttgattgtgtggggcggaatccaggacttttactttgagaaaattagagtgtttcaagcaggcacacgccttgaattcattagcatggaataataatataggacctcggttctattttgttggtttctagagctgaggtaatgattgatagggatagttgggggcattcgtatttaactgtcagaggtgaaattcttggatttgttaaagacggaatactgcgaaagcatttgccaaggatgttttca
>88e81cf401e23c4170e58b69bd9792a1eab54ef1_37
agctccaatagcgtatattaaagttgttgcggttaaaaagctcgtagttggatttctgccgaggacgaccggtccgccctctgggtgagcatctggttcggcctgggcatcttcttggagaacgtcgctgcacttgactgtgtggtgcggtatccaggacttttactttgaggaaattagagtgtttcaagcaagcacacgccttgaatacattagcatggaataataagataggacctcggttctattttgttggtttctagagctgaggtaatgattaatagggatagttgggggcattcgtatttaactgtcagaggtgaaattcttggatttgttaaagacggactactgcgaaagcatttgccaaggatgttttca
>8fdd827e5aaebff23e7141d531d4473c85c9e4b3_24
agctccagtagcgtatattaaagttgttgcggttaaaaagctcgtagttggatttctgccgaggacgaccggtccgcccactgggtgtgtatctggctcggcctgggcatcttcttggagaacgtagctgcacttgactgtgtggtgcggtatccaggacttttactttgaggaaattagagtgtttcaagcaggcgcacgccttgaatacattagcatggaataataagataggaccttggttctattttgttggtttctagagctgaggtaatgattaatagggatagttgggggcattcgtatttaactgtcagaggtgaaattcttggatttgttaaagacggactactgcgaaagcatttgccaaggatgttttca
>f15da23c46bf7d4e8ed0100e0f51d32c907361ce_33
agctccaatagcgtatattaaagttgttgcggttaaaaagctcgtagttggatttctgcttaggacgaccggtccgcccactgggtgagtatctggctcggcctgggcatcttcttggagaacgtagctgcgcttgattgtgtggtgcggtatccaggacttttactttgaggaaattagagtgtttcaagcaggcacacgccttgaatacattagcatggaataataagataggaccttggttctattttgttggtttctagagctgaggtaatgattaatagggatagttgggggcattcgtatttaactgtcagaggtgaaattcttggatttgttaaagacggactactgcgaaagcatttgccaaggatgttttca
>2efa5b52e5f4ad40b6eeda59eb49ca243500d8a5_23
agctccaatagcgtatattaatgttgttgcagttaaaaagctcgtagttggatttctgccgaggacgaccggtccgcccactgggtgtgtatctggctcggcctgggcatcttcttggagaacgtagctgcacttgactgtgtggtgcggtatccaggacttttactttgaggaaattagagtgtttcaagcaggcgcacgccttgaatacattagcatggaataataagataggacctcggttctattttgttggtttctagagctgaggtaatgattaatagggatagttgggggcattcgtatttaactgtcagaggtgaaattcttggatttgttaaagacggactactgcgaaagcatttgccaaggatgttttca
>5a6b109fba234a42b8b3dd77923a50bc85643101_22
agctccaatagcgtatattaaagttgttgcggttaaaaagctcgtagttggatttctgcttaggacgaccggtacgcccactgggtgagtatctggctcggcctgggcatattcttggagaacgtagctgcgcttgattgtgtggtgcggaatccaggacttttactttgagaaaattagagtgtttcaagcaggcacacgccttgaatacattagcatggaataataatataggacctcggttctattttgttggtttctagagctgaggtaatgattgatagggatagttgggggcattcgtatttaactgtcagaggtgaaattcttggatttgttaaagacggacaactgcgaaagcatttgccaaggatgttttca
>714697f848cfe6d73032442955fca9a8b52ba7cc_51
agctccaatagcgtatattaaagttgttgcggttaaaaagctcgtagttggatttttcaggttcagcagtacttcggtcactactgaaactggttttgacgtcatcgtcaaatagttactttgaggaaattagagtgtttaaagcagatttatgtcatgtatatattagcatggaataacaataaagattgaaagattcttgttggttacatcttttaataatgtttaatagggacaaccgggggcatttgtatttaacagtcagaggtggaattcttggatttgttaaggacaaactattgcgaaagcatttgccaaggatgttttcg
>c152628fcd8d2e43326e430ed5962bbb941ef011_21
agctccaatagcgtatattaaagttgttgcggttaaaaagctcgtagttagatttttgacctagtcgaccggtcactccaatggaatgtatcaggtttgattaggttatttgtctgatatttttggctgcacttaactgtgtggtcaaatgttcagaccttttactttgaggaaattagagtgtttcaagcaggcatttgccttgaatacgttagcatggaataataaaataagactttggttttattttgttggtttctaaaactaaagtaatgattaatagggatagtcgggggcattcgtacttaactgtcagaggtgaaattcttggatttgttaaagacgaactactgcgaaagcatttgccaaggatgttttca
>8717bf65a60a565c009620537308e2f0e57f4a16_27
agctccaatagcgtatattaaagttgttgcggttaaaaagctcgtagttggatttctgcttaggacgaccggtccgcccactgggtgagtatctggctcggcctgggcatcttcttggagaacgtagctgcgcttgattgtgtggtgcggaatccaggacttttactttgagaaaattagagtgtttcaagcaggcacacgccttgaatacattagcatggaataataatataggacctcggttctattttgttggtttctagagctgaggtaatgattgatagggatagttgggggcattcgtatttaactgtcagaggttaaattcttggatttgttaaagactgactactgcgaaagcatttgccaaggatgttttca
>a2eb76bc4cedfebc1d711ed6d6a3fba9adf1fcc5_51
agctccaatagcgtatattaaagttgttgcggttaaaaagctcgtagttggatttctgcttaggacgaccggtccgcccactgggtgagtatctggctcggcctgggcatcttcttggagaacgtagctgcgcttgattgtgtggtgcggaatccaggacttttactttgaggaaattagagtgtttcaagcaggcacacgccttgaatacattagcatggaataataagataggaccttggttctattttgttggtttctagaactgaggtaatgattaatagggatagttgggggcattcgtatttaactgtcagaggtgaaattcttggatttgttaaagacggactactgcgaaagcatttgccaaggatgttttca
>bb6c88013cfdc29d332f6e40fbbc6de3363b7ed1_23
agctccaatagcgtatattaaagttgtttcggttaaaaagctcgtagttggatttctgccgaggacgaccggtccgcccactgggtgtgtatctggctcggcctgggcatcttcttggagaacgtagctgcacttgattgtgtagtgcggtatccaggacttttactttgaggaaattaaagtgtttcaagcaggcacacgccttgaatacattagcatggaataataagataggaccttggttctattttgttggtttctagaactgaggtaatgattaatagggatagttgggggcattcgtatttaactgtcagaggtgaaattcttggatttgttaaagacggactactgcgaaagcatttgccaaggatgttttca
>c1819f88d8afe2c8ab4cf73a76cf7bc3651c1ba5_37
agctccaatagcgtatattaaagttgttgcggttaaaaagctcgtagttggatttctgctgaagatgaccggtccgccctctgggcaggtatctggctcagcttcggcatcttcttgaagaacgtctctgcacttgactgtgtggtgcggtatttaagacatttactttgaggaaattagagtgtttcaagcaagcgtgtgctttgaatacattagcatggaataataagataggacctcagtactattttgttggtttctagatctgaggtaatgattgatagggatagttgggggcattcgtatttaattgtcagaggtgaaattcttggatttgttaaagacggactactgcgaaagcatttgccaaggatgttttca
>05972c797a2c6f8d1fa32d9b5f0caf88d0b15d16_17
agctccaatagcgtatattaaagttgttgcggttaaaaagctcgtagttggatttctgcttaggactaccggtccgcccactgggtgagtatctggctcggcctgggcatcttcttggagaacgtagctgcgcttgattgtgtggggcggaatccaggacttttactttgagaaaattagagtgtttcaagcaggcacacgccttgaatacattagcatggaataataatataggacctcggttctattttgttggtttctagagctgaggtaatgattgatagggatagttgtgggcattcgtatttaactgtcagaggtgaaattcttggatttgttaaagacggactactgcgaaagcatttgccaaggatgttttca
>53d48125e33c5a57830b5c928bd94c5acc0d3e0d_50
agctctgatagtatatattaaagttgttgcggttaaaaagctcgtagttggatttctgctaaggacgaccggtccgccttcgggtgtgtacatgtgttgaccgaggcatcaatctggaagacgcctgtacttaaccgtgcgggcgtaattcagacgttttactttgaggaaattagagtgtttcaagcaggcaattgccctgaatacattagcatggaataataagataggacttcggttctattttgttggtttctagaactgaagtaatgattaatagggacagtcgggggcattcgtactcaactgtcagaggtgaaattcttggatttgttgaagacgaactactgcgaaagcatttgccaaggatgttttca
>c6d2c46f3a236eb0b657717fd2cfcee1dcf063ac_19
agctccaatagcgtatattaaagttgttgcggttaaaaagctcgtagttggatttctgccaaggacgaccggtccgccttcgggtgtgtatctgtgttgaccaaggcatctttctaggtgacgcctgtacttaactgtgtgggcgtagtctagaacatttactttgaggaaattagagtgtttcaagcaggcaattgccctgaatacattagcatggaataataagataggacttcggttctattttgttggtttctagaactgaagtaatgattaatagggacagtcgggggcattcgtactcaactgtcagaggtgaaattcttggatttgttggagacgaactactgcgaaagcatttgccaaggatgttttca
>e4143d344fd4893179ce0a22ca5bc09186a0d42e_22
agctccaatagcgtatattaaagttgttgcggttaaaaagctcgtagttggatttctgcttaggacgaccggtccgcccactgggtgagtatctggctcggcctgggcatcttcttggagaacgtagctgcgcttgattgtgtggtgcggaatccaggacttttactttgagaaacttagagtgtttcaagcaggcacacgccttgaatacattagcatggaataataatataggacctcggttctattttgttggtttctagagctgaggtaatgattgatagggatagttgggggcatttgtatttaactgtcagaggtgaaattcttggatttgttaaagactaacttatgcgaaagcatttgccaaggatgttttca
>0ca5395074e7e9058a3c5ec686d034ead835768b_21
agctctaatagcgtatattaaagttgttgcagttaaaaagctcgtagttgaatttctgctggggtagaataggtctgctctctgagtgtgtacctgatttcgaccccgcatcctctcagaatgttattggtgtactttactgtgcatcacataatatctgagatttttactttgagtaaattagagtgtttcaagcaggcatatcgccctgaatactttagcatggaataataagataggactttggttttattttattggtttatagaactaaagtaatgattaatagggatagttgggggcattcgaatttaactgtcagaggtgaaattcttagatttgttaaagtcgaactactgcgaaagcatttgccaaggatgttttca
>17a4874bc7e04edc26ec1577ae4066fc2852d455_15
agctccaatagcgtatattaaagttgttgcagttaaaaagctcgtagttgaatttctgcagaatgcggcgcgttgcacctttcgaggcgatttcgtgtttgcttctgcattctagtggctgcttagtgcacttcactgtgtactaagttgtgtgctagcttttactttgagaaaattagagtgtttcaagcaggctacttgccttgaatactgcagaatttaacttgccttgaatttaactgtcagaggtgaaattcttagatttgttaaagacgaaccaatgcgaaagcatttgccaaaaatgttttca
>6864ed4c9ab0f2a5d7a61b174eb392c47dbf7431_21
agctccaatagcgtatattaaagttgttgcggttaaaaagctcgtagttggatttctgtcaaggactaccggcccgccttcgggtgtgtgcaagtgttgaccgagacattattctgtattggcttcacttaactgtgttgtcaattttcagaatcgttactttgaggaaattagagtgtttcaagcaggcaattgccttgaatacattagcatggaataacaatataggactccggttctattttgttggtttcttgagctggagtaatgattaatagggacaatcgggggcatccgtactcaaccgtcagaggtgaaattcttggattggttgaagacgaactactgcgaaagcatttgccaaggatgttttca
>7281a300e2fbacbfb132bc33179513ae99bc9a2f_21
agctccaatagcgtatattaaagttgttgcggttaaaaagctcgtagttggatttctgccgaggacgaccgatcgacccatacgggtttgtatctggtttgaccgaggcatcctcttggaatacgtatagcactttactgggttatgcggagtccaagacttttactttgaggaaattagagtgtttcaagcaggcatttgccgtgaatacattagcatggaataataagataggactttggttctattttgttggtttctggaactgaagtaatgattaatagggacagtcgggggcattcgtacttaactgtcagaggtgaaattcttggatttgttaaagacgaactactgcgaaagcatttgccaaggatgttttca
>87a9a37400a625a85e0f1bba40d2834aa50f42e5_61
agctccaatagcgtatattaaagttgttgcggttaaaaagctcgtagttggatttctgctgaggacgaccggtccgccccctgggtgagcatctggttcggcctgagcatcttcttggagaatgttgctgcacttcactgtgtggcgcggtatccaggacttttactttgaggaaattagagtgcttcaagcaggcacacgccttgaacacgtgagcatggaataatgtgataggacctcggttctattttgttggtttctagagctgaggtaatgataaatagggacagttgggggcattcgtatttaactgtcagaggtgaaattcttggatttgttaaagacggaccactgcgaaagcatttgccacggatgtcttca
>00bc87dcd8b04c7d2f678d44d7ffe3fb072b60c2_23
agctccaatagcgtatattaaagttgttgcagttaaaaagctcgtagttggatttctgttcaggatgacgggtccaatgtttcattgtgtatctgtgcagcctgagcattttcttggggaacgtcactgcccttaattgtgtggtgcggtatccaggaccgttactttgagaaaattagagtgtttcaagcaagcctacgctttgaatactgcagcatggaataatatgataagacttcggttttattttgtgggtttctaaaactgaagtgatgattaatagggacagttgggggcattcgtatttaactgtcagaggtgaaattcttggatttgttaaagacgaactactgcgaaagcatttgccaaggatgttttca
>4d43cbbc7f7eda599d52b0cfb45eb9e680e5f73f_19
agctccaatagcgtatattaaagttgttgcggttaaaaagctcgtagttggatttctgcttaggacgaccggtccgccaactgggtgagtatctggctcggcctgggcatcttcttggagaacgtagctgcgcttgattgtgtggggcggaatccaggacttttactttgagaaaattagagtgtttcaagcaggcacacgccttgaatacattagcatggaataataatataggacctcggttctattttgttggtttctagagctgaggtaatgattgatagggatagttgggggcattcgtatttagttgtcagaggtgaaattcttggatttgttaaagacggactactgcgaaagcatttgccaaggatgttttca
>77d331b6c643673e276b62b2e15516237f8ccc95_20
agctccaatagcgtatattaaagttgttgcggttaaaaagctcgtagttggatttctgccgaggacgaccggtccgccctctgggtgagcatctggttcggcctgggcatcttcttggagaacattgctgcacttgactgtgtggtgcggtatccaggacttttactttgaggaaattagaatgtttcaagcaggcacacgccttgaatacattagcatggaataataagataggacctcggttctattttgttggtttctagagctgaggtaatgattaatagggatagttgggggcattcgtatttaactgtcagaggtgaaattcttggattgttaaagacggactactgcgaaagcatttgccaaggatgttttca
>d089ac883663426a904bc9f2931c308f192daf21_17
agctccaatagcgtatattaaagttgttgcggttaaaaagctcgtagttggatttctgcttaggacgaccggtccgcccactgggtgagtatctggctcggcctgggcatcttcttggagaacgtagctgcgcttaattgtgtggggcggaatccaggacatttactttgagaaaattagagtgtttcaagcaggcacacgccttgaatacattagcatggaataataatataggacctcggttctattttgttggtttctagagctgaggtaatgattgatagggatagttgggggcattcgtatttaactgtcagaggtgaaattcttggatttgttaaagacggactactgcgaaaccatttgccaaggatgttttca
>e15cb9898273060c229841f7fb78909e71605eda_26
agctccaatagcgtatattaaagttgttgcggttaaaaagctcgtagttggatttctgccgaggacgaccggtccgccctctgggtgagtatctggcttggcctgggcatcttcttggagaacgtagctgcacttgactgtgtggtgcggtatccaggacttttactttgaggaaattagagtgtttcaagcaggcacacgccttgaatacattagcatggaataataagataggaccttggttctattttgttggtttctagagctgaggtaatgattaatagggatagttgggggcattcgtatttaactgtcagaggtgaaattcttggatttgttaaagacggactactgcgaaagcatttgccaaggatgttttca
>052e375b1b8b03de9eabd6a56c290fd8bf2bdce8_21
agctccaatagcgtatattaaatttgttgcggttaaaaagctcgtagttggatttctgccgaggacgaccggtccgcccactgggtgtgtatctggctcggcctgggcatcttcttggagaacgtagctgcacttgactgtgtggtgcggtatccaggacttttactttgaggaaattagagtgtttcaagcaggcacacgccttgaatacattagcatggaataataagataggacctcggttctattttgttggtttctagagctgaggtaatgattaatagggatagttgggggcattcgtatttaactgtcagaggtgaaattcttggatttgttaaagacggactactgcgaaagcatttgccaaggatgttttca
>3f0e54d8c01430218d451533f9f0a51a73c940c2_14
agctccaatagcgtatattaaagttgttgcggttaaaaagctcgtagttggatttctgtcgaggacgatcggtccgcctcctgggtgagtatctggctcggccttggcatcttcttggagaacgtactgcacttcattgtgtggtgcggaatccaggacctttactttgaggaaattagagtgtttcaagcaggcgcacgccttgaatacattagcatggaataataagataggacctcggttctattttgttggtttctagagctgaggtaatgattaatagggatagttgggggcattcgtatttaactgtcagaggtgaaattcttggatttgttaaagacggactactgcgaaagcatttgccaaggatgttttca
>6081f75ff97eecd031bcbf61649012efaf8bdb9c_37
agctctgatagtatatattaaagttgttgcggttaaaaagctcgtagttggatttctgccgaggacgaccggtccgcccactgggtgtgtatctggctcggcctgggcatcttcttggagaacgtagctgcacttgactgtgtggtgcggtatccaggacttttactttgaggaaattagagtgtttcaagcaggcacacgccttgaatacattagcatggaataataagataggaccttggttctattttgttggtttctagaactgaggtaatgattaatagggatagttgggggcattcgtatttaactgtcagaggtgaaattcttggatttgttaaagacggactactgcgaaagcatttgccaaggatgttttca
>8aedb764b91c410e79af9a410626e2d190432588_18
agctccaatagcgtatattaaagttgttgcggttgaaaagctcgtagttggatttctgccgaggacgaccggtccgccctctgggtgagcatctggttcggcctgggcatcttcttggagaacgtcgctgcacttgactgtgtggtgcggtatccaggacttttactttgaggaaattagagtgtttcaagcaggcacacgccttgaatacattagcatggaataataagataggacctcggttctattttgttggtttctagagctgaggtaatgattaatagggatagttgggggcattcgtatttaactgtcagaggtgaaattcttggatttgttaaagacggactactgcgaaagcatttgccaaggatgttttca
>a3f25554630d8b611b934cd5129e486ee8f74c45_16
agctccaatagcgtatattaaagttgttgcagttaaaaagctcgtagttggatttctgtttaggatgaccggtccactttcgggtggttacctggtttagcttaaacattttcttggagaacgttgctgcacttaattgtgtggtgcggaatccaggacttttactttgagaaaattagagtgtttcaagcaggccaacgctttgaatactacagcatggaataatattataagacttcggttttattttgtgggtttctaaaactgaagtaatgattaatagggacagttgggggcattcgtatttaactgtcagaggtgaaattcttggatttgttaaagacgaactactgcgaaagcatttgccaaggatgttttca
>a9b2531cf2d5178e21d4aaa7dcc622e7606cce16_14
agctccaatagcgtatattaaagttgttgcggttaaaaagctcgtagttggatttctgccaaggacgaccggtccgcccactgggtgtgtatctggctcggcctgggcatcttcttggagaacgtagctgcacttgactgtgtggtgcggtatccaggacttttactttgaggaaattagagtgtttcaagcaggcacacgccttgaatacattagcatggaataataagataggaccttggttctattttgttggtttctagaactgaggtaatgattaatagggatagttgggggcattcgtatttaactgtcagaggtgaaattcttggatttgttaaagacgaactactgcgaaagcatttgccaaggatgttttca
>05a727ecc8072d8ff4e676fe70250eeb661e2aa5_11
agctccaatagcgtatattaaagttcttgcggttaaaaagctcgtagttggatttctgccgaggacgaccggtccgcccactgggtgtgtatctggctcggcctgggcatcttcttggagaacgtatctgcgcttgactgtgtggtgcggcatccaggacttttactttgaggaaattagagtgtttcaagcaggcacatgccttgaatacattagcatggaataataagataggaccttggttctgttttgttggtttctagagctgaggtaatgattaatagggatagttgggggcattcgaatttaactgtcagaggtgaaattcttggatttgttaaagacgaactactgcgaaagcatttgccaaggatgttttcc
>223ed0fe0e2696e07d5888992182926f8cdb6d2f_11
agctccaatagcgtatattaaagttgttgcggttaaaaagctcgtagttggatttctgccaaggacgaccggtccaccttagggtgtgtatctgtgttgaccgaggcatctttctaggtgacgcccgtgcttcactgtgcgacgcgtgctctagaacttttactttgaggaaattagagtgttccaagcaggcttatgccctgaatacattagcatggaataataaggtaggacttcggttctattttgttggtttctagaactgaagtaatgattaatagggacagtcgggggcattcgtactcaactgtcagaggtgaaattcttggatttgttggagacgaactagctactgcgaaagcatttgccaaggatgttttca
>2c7f5ae8e54dc84205013a4ace2b3f486aee8903_13
agctctgatagtatatattaaagttgttgcggttaaaaagctcgtagtcgctcttctgtcagaaacgaccggtccgccctttgggtgagtatctggctcggctctagcctcttcttggttgaccatgctgcacttgagtgtgtggcaagggacccaggactttcactttgagaaaattggagtgtttcaagcaggctcacgccgtggacacataagcatggaatgaatggtttgaacctcgcgtctatcttgttggtttctagaaatggggtgatcaagatagggatagttgggggtattcgtatttaactgtcagaggtgaaattcttggatttgttaaagacggaccactgcgaaagcatttgccaaagatgttctca
>3d184ea6a8005e325a98496bc2c0e46d90a0b570_23
agctccaatagcgtatattaaagttgttgcagttaaaaagctcgtagttgaatttctgctggagttgaataggtctgctctttgagtgtgtaccttgtttcgacttcgcatcctcttagaatgttattaaagtactttattgtgctttatataatatctaagatttttactttgagtaaattagagtgtttcaagcaggcatattgccttgaatactttagcatggaataataaaataggattttggttctattttattggtttatagaactaaagtaatgattaatagggacagttgggggcattcgaatttaactgtcagaggtgaaattcttagatttgttaaagtcgaactactgcgaaagcatttgccaagaatgttccca
>478a288b1bd9efce4a7784f481579015b448dfea_10
agctccaatagcgtatattaaagttgttgcagttaaaaagctcgtagttggatttctgttcaggatgaccggtccacttcggtggtcacctggtttagcctgaacattttcctggagaacgtcactgcacttcattgtgtggtgtggaatccaggacctttactttgagaaaattagagtgtttcaagcaggccaacgctttgaatactgcagcatggaataatatgataagacttcggttttattttgtgggtttctaaaactgaagtaatgattaatagggacagttgggggcatttgtatttaactgtcagaggtgaaattcttggatttgttaaagacaaactactgcgaaagcatttgccaaggatgttttca
>4cfa51b52bf80fb0132dbeee7c77311d55ebbbc3_14
agctccaatagcgtatattaaagttgttgcggttaaaaagctcttagttggatttctgccgaggacgaccggtccgcccactgggtgtgtatctggctcggcctgggcatcttcttggagaacgtagctgcacttgactgtgtggtgcggtatccaggacttttactttgaggaaattagagtgtttcaagcaggcacacgccttgaatacattagcatggaataataagataggaccttggttctattttgttggtttctagagctgaggtaatgattaatagggatagttgggggcattcgtatttaactgtcagaggtgaaattcttggatttttggaagacggactactgcgaaagcatttgccaaggatgttttca
>4db83ae1a5b02c7637ee985a00935ea6da0034a9_10
agctccaggagcgtatattaaagttgttgcggttaaaaagctcgtagttggatttctgctgaggacgaccggtccgccctctgggtgagcatctggctcggcctgagcatcctcttggagaaggtgtgtgcactttactgtgtgcacccgtatccaggacttttactttgaggaaattggagtgcttcaagcaggcacacgccttgaacacatgagcatggaataatgcgataggacctcggttctattttgttggtttctagaactgaggtaatgataaatagggacagttgggggcattcgtatttaactgtcagaggtgaaattcttggatttgttaaagacggaccactgcgaaagcatttgccacggatgtcttca
>92e8d7273417b4283b07356893bcd9335a41966a_23
agctccaatagcgtatattaaagttgttgcggttaaaaagctcgtagttggatttctgcttaggacgaccggtccgcccactgggtgagtatctggctcggcctgggcatcttcttggagaacgtagctgcgcttgattgtgtggtgcggaatccaggacttttactttgaggaaattagagtgtttcaagcaggcgcacgccttgaatacattagcatggaataataagataggacctcggttctattttgttggtttctagagctgaggtaatgattaatagggatagttgggggcattcgtatttaactgtcagaggtgaaattcttggatttgttaaagacggactactgcgaaagcatttgccaaggatgttttca
>f590b48a342ddad6a9b77178cd18445c05376fd3_13
agctccaatagcgtatattaaagttgttgcggttaaaaagctcgtagttggatttctgcttaggacgaccggtccgcccactgggtgagtatctggctcggccttggcatcttcttggagaacgtagctgcgcttgattgtgtggggcggaatccaggacttttactttgagaaaattagagtgtttcaagcaggcacacgccttgaatacattagcatggaataataatataggacctcggttctattttgttggtttctagagctgaggtaatgattgatatggatagttgggggcattcgtatttaactgtcagaggtgaaattcttggatttgttaaagacggactactgcgaaagcatttgccaaggatgttttca
>0201701ea9ed0304fa8260cdc5f8b790895c17d6_14
agctccaatagcgtatattaaagttgttgcggttaaaaagctcgtagttggatttctgctgaggacgatcggtccgccctctgggtgagtatctggctcggccttggcatcttcttggagaacgtagctgcacttgactgtgtggcgcggtatccaggacttttactttgaggaaattagagtgtttcaagcaggcgcacgccttgaatacattagcatggaataataagataggaccttggttctattttgttggtttctagaactgaggtaatgattaatagggatagttgggggcattcgtatttaactgtcagaggtgaaattcttggatttgttaaagacggactactgcgaaagcatttgccaaggatgttttca
>8c5a62d7716b4a5bfe911645a800d36e59afe6dd_18
agctccaggagcgtatattaaagttgttgcggttaaaaagctcgtagttggatttctgttgaggacgatcggtccgccttctgggtgagtatctggctcggccttggcatcttcttggagaacgtactgcacttcattgtgtggtgcggaatccaggacctttactttgaggaaattagagtgtttcaagcaggcgcacgccttgaatacattagcatggaataataagataggacctcggttctattttgttggtttctagagctgaggtaatgattaatagggatagttgggggcattcgtatttaactgtcagaggtgaaattcttggatttgttaaagacggactactgcgaaagcatttgccaaggatgttttca
>29deeb67c772ac4044ec71744905921a9f68ee5c_9
agctccaatagcgtatattaaagctgttgcggttaaaaagctcgtagttggacttctgctgaggacgactggtctgccctcagggtgagtatctggttcggccgaagcatcttcttggaggttgcaagcgcacttcactgtgtggtgcaaccgactggaattttactttgaggaaattagagtgttcaagggaggcgtacgcctcgcataggttagcatggaataatgctgtaagaccatggtattgctgtgttggtttctatcacctaaggtaatgataaacagggatagttgggggcattcgtatttaacagtcagaggtgaaattcttggatttgttaaagacgaacgactgcgaaagcatttgccaaggatgttttca
>44f11409dfc8e3a951af2ffe4c26da1f8b26ba60_26
agctccaatagcgtatattaaagttgttgcggttaaaaagctcgtagttggatttctgctgaggacgaccggtccgccctctgggtgagtatctggctcggccttggcatcttcttggagaacgttactgcacttgattgtgtggtgcggtatccaggacttttactttgaggaaattagagtgtttcaagcaggcacacgccttgaatacattagcatggaataataagataggacctcggttctattttgttggtttctagagctgaggtaatgattaatagggatagttgggggcattcgtatttaactgtcagaggtgaaattcttggatttgttaaagacggactactgcgaaagcatttgccaaggatgttttca
>4b8ba3c5e3cb0ebb0c0353e28b73998824a941fc_8
agctccaatagcgtatattaaagttgttgctgttaaaaagctcgtagttggatttctgcttaggacgaccggtccgcccactgggtgagtatctggctcggcctgggcatcttcttggagaacgtagctgcgcttgattgtgtggtgctgaatccaggacttttactttgagaaaattagagtgtttcaagcaggcacacgccttgaatacattagcatggaataataatataggacctcggttctattttgttggtttctagagctgaggtaatgattgatagggatagttgggggcattcgtatttaactgtcagaggtgaaattcttggatttgttaaagacggactactgcgaaagcatttgccaaggatgttttca
>569d0c5bc6ed01e1b559e01eed5685ae66ab2228_12
agctccaatagcgtatattaaagttgttgcggttaaaaagctcgtagttggatttctgctgaggacgatcggtccgccctctgggtgagtatctggctcggccttggcatcttcttggagaacgttgctgcacttgactgtgtggcgcggtatccaggacttttactttgaggaaattagagtgtttcaagcaggcgcacgccttgaatacattagcatggaataataagataggaccttggttctattttgttggtttctagagctgaggtaatgattaatagggatagttgggggcattcgtatttaactgtcagaggtgaaattcttggatttgttaaagacggactactgcgaaagcatttgccaaggatgttttca
>64ec4a367b3035082ade2b723ec9758927bdded8_17
agctccaggagcgtatattaaagttgttgcggttaaaaagctcgtagttggattttatagtcgaagctacattttggtgtaactcgattatggtaataatttattattacacgttactttgaggaaattagagtgtttaaagcagatctttgtcatgtatatattagcatggaataacactaaggattattaaaactttgttggttactttttttaataatgattgatagggacagccgggggcatttgtatttaacagtcagaggtggaattcttggatttgttaaggacaaactattgcgaaagcatttgccaaggatgttttcg
>7e42418129ea5aaea8d3149fba519f32ca6ad996_26
agctccaatagcgtatattaaagttgttgcggttaaaaagctcgtagttggatttctgcttaggacgaccggtccgcccactgggtgagtatctggctcggcctgggcatcttcttggagaacgtagctgcgcttgattgtgtggtgcggaatccaggacttttactttgagaaaattagagtgtttcaagcaggcacacgccttgaatacattagcatggaataataatataggacctcggttctattttgttggtttctagagctgaggtaatgattgatagggatagttgggggcattcgtatttaactgtcagaggtgaaattcttggatttgttaaagactaacttatgcgaaagcatttgccaaggatgttttca
>84e420408460fd0519b7dc846f8239a6e1aba082_9
agctccaatagcgtatattaaagttgttgcggttaaaaagctcgtagttggatttctgcttaggacgaccggtccgcccttgagcatctggctcggcctgggcatcttcttggagaacgtagctgcgcttgattgtgtggtgcggaatccaggacttttactttgagaaacttagagtgtttcaagcaggcacacgccttgaatacattagcatggaataataatataggacctcggttctattttgttggtttctagagctgaggtaatgattgatagggatagttgggggcattcgtatttaactgtcagaggtgaaattcttggatttgttaaagacggactactgcgaaagcatatgccaaggatgttttca
>a2357d1f3dd7275476e8e5d782deeab3038aa403_13
agctccaatagcgtatattaaagttgttgcagttaaaaagctcgtagttgaatttctgcgggagttgaataggtctgctctttgagtgtgtaccttatttcgacttcgcatcctcttagaagaatagtaaagtactttattgtgctttatatataatctaagacctttactttgagtaaattagagtgtttcaagcaggcatattgccttgaatactttagcatggaataataaaataggactttggttctattttattggtttatagaactaaagtaatgattaatagggacagttgggggcattcgaatttaactgtcagaggtgaaattcttagatttgttaaagtcgaactactgcgaaagcatttgccaaggatgttttca
>35ec737728ca4e6f498a12719de09b79099c27d5_12
agctcgatgaacgaacacgatctttgttgcagttaaaaagctcgtagttggacttctgctgggcaccaagccaagaaagggcaacctttctaggtcatggttgcccagcatgctcgtgagtgggtcactccttgattggagtgtgccagggctcacgtctgttactgtgaacaaaacggagcgttcatagcaggtttttggcctgaacgactcgcacgggataactcaaatggactcttgttcatttcgttggtttggaatgagggtttggttaaaaggaacaatggggggtatttgtactgtcacgctagaggtgaaattcttggattgtgacatgacaaacgactgcgaaagcatttaccaaaaatgttttca
>ef02f2a97f01c3be157c52e25513b9477856f56c_6
agctccaatagcgtatattaaagttgttgcggttaaaaagctcgtagttggatttctgcttaggacgaccggtccgcccactgggtgagtatctggctcggcctgggcatcttcttggagaacgtagctgcgcttgattgtgtggtgcggaatccaggacttttactttgagaaaattagagtgtttcaagcaggcacacgccttgaatacattagcatggaataataatataggacctcggttctattttgttggtttctagagctgatgtaatgattgatagggatagttgggggcattcgcatttaactgtcagaggtgaaattcttggatttgttaaagacggactactgcgaaagcatatgccaaggatgttttca
>0b32ab63926c0405c1441d7fed5db87a70cdcc79_9
agctccaatagcatatattaaagttgttgcagttaaaaagctcgtagttggatttctgttcaggatgaccggtccaatttcggttgtgtacctgggatagcctgaacattttcctggagaacgtcactgcacttgattgtgtggtgcggtatccaggacctttactttgagaaaattagagtgtttcaagcaagcctacgctttgaatactgcagcatggaataatatgataagacttcggttttattttgtgggtttctaaaactgaagtaatgattaatagggacagttgggggcattcgtatttaactgtcagaggtgaaattcttggatttgttaaagacgaactactgcgaaagcatttgccaaggatgttttca
>0e12e913708a090ec2aa02d10086882f87c721b3_15
agctccaatagcgtatattaaagttgttgcggttaaaaagctcgtagttggatttctgctaagggctaccggtccgccttcgggtgtgaatcagtgtggtccaaagcatctgtctagtggcgctcatgcttaactgtgtgatgcgtattctagaccttttactttgaggaaattagagtgtttcaagcaggcatatgccgtgaatacattagcatggaataataatttaggacttcggtcctattttgttggtttctggactgaagtaatgattaataggaacaatcgggggcattcgtattcaacagtcagaggtgaaattcttggatttgttggagacgaactactgcgaaagcatttgccaaggatgttttca
>7f82fcfab420aadea863c2d4fe7fd6b65aefd4ba_5
agctccaatagcgtatattaaagttgttgcagttaaaaagctcgtagttgaatttctgcagaatgcggcgcgatgcaccttttaggcgatttcgtgtttgcttctgcattctagtgactgctgtatacactttactgtgtataccgtttttcactagcatttactttgagaaaattagagtgtttcaagcaggctttttgccttgaactgaagtaatgattaacagggacagttgggggcatttgaatttaactgtcagaggtgaaattcttagatttgttaaagacgaaccaatgcgaaagcatttgccaaaaatgttttca
>c50196f46f1ce669b41fb6cf7f2782521bf1faba_8
agctccaggagcgtatattaaagttgttgcggttaaaaagctcgtagttggatttctgtcgaggacgaccggtccgccctccgggtgagtatctggttcggcctaggcatcctcttggagaaggagcggtcactttgctgtgaacgcccgtatccaggacttttactttgaggaaattggagtgcttcaagcaggcacacgccttgaacacatgagcatggaataatgcgataggacctcggttctattttgttggtttctagagctgaggtaatgataaatagggacagttgggggcattcgtatttaactgtcagaggtgaaattcttggatttgttaaagacggaccactgcgaaagcatttgccacggatgtcttca
>d112d3d56666affd015d5908211b0a54b14fe6ce_5
agctccaatagcgtatattaaagttgttgcggttaaaaagctcgtagttggatttctgctgaggacgaccggtccgccctctgggtgagcatctggctcggcctgagcatcctcttggagaaggtgtgtgcactttactgtgtgcacccgtatccaggacttttactttgaggaaattggagtgcttcaagcaggcacacgccttgaacacatgagcatggaataatgcgataggacctcggttctattttgttggtttctagaactgaggtaatgataaatagggacagttgggggcattcgtatttaactgtcagaggtgaaattcttggatttgttaaagacggaccactgcgaaagcatctgccaagtacgttccca
>04ef7824b6bc77c4bd755fa360e2903b5a018a47_8
agctccaatagcgtatattaaagttgttgcggttaaaaagctcgtagttggatttctgcttaggacgaccggtccgcccactgggtgagtatctggctcggcctgggcatcttcttggagaacgtagctgcacttgactgtgtggtgcggtatccaggacttttactttgaggaaattagagtgtttcaagcaggcacacgccttgaatacattagcatggaataataagataggaccttggttctattttgttggtttctagaactgaggtaatgattaatagggatagttgggggcattcgtatttaactgtcagaggtgaaattcttggatttgttaaagacggactactgcgaaagcatttgccaaggatgttttca
>1d6727aeab4b54c40b50810aa80e17a32ef0dd19_12
agctccaatagcgtatattaaagttgttgcagttaaaaagctcgtagttggatttctgcctaggacgaccggtccgcccactgggtgagtatctgactcggcctgggcatcttcttggagaacgtagctgcgcttgattgtgtggtgcggaatccaggacttttactttgagaaaattagagtgtttcaagcaggcacacgccttgaatacattagcatggaataataatataggacctcggttctattttgttggtttctagagctgaggtaatgattgatagggatagttgggggcattcgtatttaactgtcagaggtgaaattcttggatttgttaaagacggactactgcgaaagcatttgccaaggatgttttca
>2e7729e1d1a799dd75d5d1cc19cd093e6eb128a4_4
agctccaatagcgtatattaaagttgttgcggttaaaaagctcgtagttggatttctgcttaggacgaccggtccgcccactgggtgagtatctggctcggcccgggcatcttcttggagaacgtagctgcgcttgattgtgtggtgcggaatccaggacttttactttgagaaaattagagtgtttcaagcaggcacacgccttgaatacattagcatggaatcataatataggacctcggttctattttgttggtttctagagctgaggtaatgattgatagggatagttgggggcattcgtatttaactgtcagaggtgaaattcttggatttgttaaagacggactactgcgaaagcatttgccaaggatgttttca
>351b1077e3e4b89d87dcdfc9ec20a475ea190eb7_22
agctccaatagcgtatattaaagttgttgcggttaaaaagctcgtagttggatttctgccgaggacgaccggtccgcccactgggtgtgtatctggctcggcctgggcatcttcttggaggacgtatctgcacttgactgtgtggtgcggcatccaggacttttactttgaggaaattagagtgtttcaagcaggcacatgccttgaatacattagcatggaataataaggtaggaccttggttctattttgttggtttctagaactgaggtaatgattaatagggatagttgggggcattcgaatttaactgtcagaggtgaaattcttggatttattaaagacgaactactgcgaaagcatttgccaaggatgttttcg
>44cb591706f5d8e6a8f94fb23b187c60f93fc502_4
agctctcaaagtgtatatcgtcattgctgcggttaaaaagctcgtagttggatttctgctgaggacgaccggtccgccctctgggtgagtatctggctcggccttggcatcttcttggagaacgttactgcacttgattgtgtggtgcggtatccaggacttttactttgaggaaattagagtgtttcaagcaggcacacgccttgaatacattagcatggaataataagataggaccttggttctattttgttggtttctagaactgaggtaatgattaatagggatagttgggggcattcgtatttaactgtcagaggtgaaattcttggatttgttaaagacggactactgcgaaagcatttgccaaggatgttttca
>5eed2099d239d05fd3f578af04d0a8eb81719fe0_9
agctccaatagcgtatattaaagttgttgcggttaaaaagctcgtagttggatttctgcttaggacgaccggtccgcccactgggtgagtatctggctcggcctgggcatcttcttggagaacgtagctgcgcttgattgtgtggtgcggaatccaggacttttactttgaggaaattagagtgtttcaagcaggcatacgccttgaatacattagcatggaataataagataggacctcggttctattttgttggtttctagatctgaggtaatggttaatagggatagttgggggcattcgtatttaactgtcagaggtgaaattcttggatttgttaaagacggactactgcgaaagcatttgccaaggatgttttca
>7551555993b7191609cab1b611545d5ac858283b_23
agctccaatagcgtatattaaagttgttgcggttaaaaagctcgtagttggatttctgccgaggacgaccggtccgccctctgggtgagtatctggctcggcctgggcatcttcttggagaacgtatctgcacttgactgtgtggtgcggtattcaggacttttactttgaggaaattagagtgtttcaagcaggcatacgccttgaatacattagcatggaataataagataggacctcggttctattttgttggtttctagatctgaggtaatgattgatagggatagttgggggcattcgtatttaactgtcagaggtgaaattcttggatttgttaaagacggactactgcgaaagcatttgccaaggatgttttca
>a54f54dd3bfac045239dea1ce480d26dc5beac41_6
agctccaatagcgtatattaaagttgttgcagttaaaaagctcgtagttggatttctggttagaccgatcgacccgcccattgggtgtgcgtctggtcgagtctatccatccttctggagaacgtcgctgtcattcacttggtggcggcggtatccaggacttttactttgaaaaaattagagtgtttaaagcaggccaacgctttgaatacattagcatggaataataaaataggactacgtttctattttgttggtttctaggaacgcagtaatgattaatagggatagttgggggcattcgtatttaatagtcagaggtgaaattcttggatttgttaaagacgaactactgcgaaagcatttgccaaggatgttttca
>b08bcb283d7095325bdb4399e853e6aea7bd30df_7
agctccaatagcgtatattaaagttgttgcggttaaaaagctcgtagttggatttctgccgaggacgaccggtccgccctctgggtgagtatctggctctgcctgggcatcttcttggagaacgtagctgcacttcactgtgtggtgcggtatccaggacttttactttgaggaaattagagtgtttcaagcaggcatacgccttgaatacattagcatggaataataagataggacctcggttctattttgttggtttctagatctgaggtaatggttaatagggatagttgggggcattcgtatttaactgtcagaggtgaaattcttggatttgttaaagacggactactgcgaaagcatttgccaaggatgttttca
>c6abc78693ef3bcc644f9e819d7512fbed1ab6f8_5
agctctgatagtatatattaaagttgttgcggttaaaaagctcgtagttggatttctgcttaggacgaccggtccgcccactgggtgagtatctggctcggcctgggcatcttcttggagaacacaactgcacttgactgtgtggtgtggtatccaggacttttactttgagaaaattagagtgtttcaagcaggcacacgccttgaatacattagcatggaataataatataggacctcggttctattttgttggtttctagagctgaggtaatgattgatagggatagttgggggcattcgtatttaactgtcagaggtgaaattcttggatttgttaaagacggactactgcgaaagcatttgccaaggatgttttca
>c8f36d9238137375044edf00da8cad84ab231937_10
agctccaaaagcgtatattaaagttgttgcggttaaaaagctcgtagttggattttatagtcgaagctacattttggtgtaactcgattatggtaataatttattattacacgttactttgaggaaattagagtgtttaaagcagatctttgtcatgtatatattagcatggaataacactaaggattattaaaactttgttggttactttttttaataatgattgatagggacagccgggggcatttgtatttaacagtcagaggtggaattcttggatttgttaaggacaaactattgcgaaagcatttgccaaggatgttccca
>d5f4ab857f9cd02ba17760a079c979c29f4a826d_7
agctccaggagcgtatattaaagttgttgcggttaaaaagctcgtagttggatttctgctgaggacgaccggtccgccctccgggtgagcatctggctcggcctttgcatcttcttggggaacgttgctgcacttcactgtgtggcgcggtatccaggacttttactttgaggaaattagagtgcttcaagcaggcacacgccttgaacacgtgagcatggaataatgtgataggacctcggttctattttgttggtttctagagctgaggtaatgataaatagggacagttgggggcattcgtatttaactgtcagaggtgaaattcttggatttgttaaagacggaccactgcgaaagcatttgccacggatgtcttca
>edf646d1ce9e460020299f0cce2f2371dfaf8ffc_5
agctccaatagcgtatatttaagttgttgcagttaaaaagctcgtagttggatttctgccgaggacgaccggtccgcccactgggtgtgtatctggctcggcctgggcatcttcttggagaacgtagctgcacttgactgtgtggtgcggtatccaggacttttactttgaggaaattagagtgtttcaagcaggcacacgccttgaatacattagcatggaataataagataggacctcggttctattttgttggtttctagagctgaggtaatgattaatagggatagttgggggcattcgtatttaactgtcagaggtgaaattcttggatttgttaaagacggactactgcgaaagcatttgccaaggatgttttca
>027bbb34b140df15b28de6788e154eb7a3c25256_8
agctccaagagcatatattaaagttgttgcagttaaaaagctcgtagttggatttctgcttaggacgaccggtccgcccactgggtgagtatctggctcggcctgggcatcttcttggagaacgtagctgcgcttgattgtgtggtgcggaatccaggacttttactttgagaaaattagagtgtttcaagcaggcacacgccttgaatacattagcatggaataataatataggacctcggttctattttgttggtttctagagctgaggtaatgattgatagggatagttgggggcattcgtatttaactgtcagaggtgaaattcttggatttgttaaagacggactactgcgaaagcatttgccaaggatgttttca
>05eef0c777d38968907e549ac5fda0fae976569e_11
agctccaagagcgtagagtaaagttgttgcggttaaaaagctcgtagttggatttctgcttaggacgaccggtccgcccactgggtgagtatctggctcggcctgggcatcttcttggagaacgtagctgcgcttgattgtgtggtgcggaatccaggacttttactttgagaaaattagagtgtttcaagcaggcacacgccttgaatacattagcatggaataataatataggacctcggttctattttgttggtttctagagctgaggtaatgattgatagggatagttgggggcattcgtatttaactgtcagaggtgaaattcttggatttgttaaagacggactactgcgaaagcatttgccaaggatgttttca
>08d0ee660179cbc23a42c5cac56223344ddddaa5_7
agctccaatagcgtatattaaagttgttgcagttaaaaagctcgtagttgaatttctgcgggagttgaataggtctgctctttgagtgtgtaccttatttcgacttcgcatcctcttagaatattattaaagtactttattgtgctttatataatatctaagatctttactttgagtaaattagagtgtttcaagcaggcatcttgccttgaatactttagcatggaataataaaataggactttggttctattttattggtttatagaactaaagtaatgattaatagggacagttgggggcattcgaatttaactgtcagaggtgaaattcttagatttgttaaagtcgaactactgcgaaagcatttgccaaggatgttttca
>219b58b56075653d073f23e624f1231d2d51b2a9_4
agctccaatagcgtatattaaagttgttgcggttaaaaagctcgtagttggatttctgcttaggatgaccggtccgccttcgggtgtgtatcagcgttgtccagagcatctttctagttgcgcccgtgcttcaatgtacggtgcgtgttctagatcttttactttgaggaaattagagtgtttcaagcaggcatatgccttgaatactttagcatggaataataagataggacttcggttctattttgttggtttctagaactgaagtaatgattaatagggacaatcgggggcattcgtactcaatagtcagaggtgaaattcttggatttgttggagacgaactactgcgaaagcatttgccgaggatgttttca
>25c273820cb13e883701fe04d21f30671394dfdd_4
agctccaagagcatatattaaagttgttgcagttaaaaagctcgtagttggatttctgctgaggatgaccggtccgccctctgggtgagtatctggctcagccttggcatcttcctgaagaacgttgctgcacttgactgtgtggtgcggtatttaggacatttactttgaggaaattagagtgtttcaagcaagcgcacgccttgaatacattagcatggaataataagataggacctcggttctattttgttggtttctagagctgaggtaatgattgatagggatagttgggggcattcgtatttaactgtcagaggtgaaattcttggatttgttaaagacggactactgcgaaagcatttgccaaggatgttttca
>34f4882d7a687b1704629152f3160c66a50bb507_4
agctccaatagcgtatattaaagttgttgcggttaaaaagctcgtagttggatttctgccgaggacgaccggtccgcccactgggtgtgtatctggctcggcctgggcatcttcttggagaacgtagctgcacttgactgtgtggtgcggtatttaggacatttactttgaggaaattagagtgtttcaagcaagcgcacgccttgaatacattagcatggaataataagataggacctcggctctatttcgttggtttctagagctgaggtaatgattgatagggacagttgggggtattcgtattccattgtcagaggtgaaattcttggatttactgaagactaactactgcgaaagcatttgccaaggacgttttca
>375232b46264968ca639d68b54706febd5940eef_7
agctccaatagcgtatattaaagttgttgcagttaaaaagctcgtagtcgaaagtctgcgggggcggggggcgtgtcacttctggtgatgcgcgcccacagagatatcgtaagagtctcggcccctgcattgggagcggcgctgtgctgggtttcatcgcccggcacggattgtgcgctccgagttactttgaggaaaatagagtgttcaaagcaggcaatgcagcttgaatatttatgcatggaataacaggacaaggcgggccgcctatgtttgttggtgacctccggggagacgcgtcatggtgaatagggacagttgggggcattcatattcagaagctagaggtgaaattcttggattttctgaagatggactagtgcgaaagcatttgccaaggatgttttca
>3c69faa7cccdb765a46ca2cc9a11335d17a37c5b_3
agctccagtagcgtatattaaagttgttgcagttaaaaagctcgtagttggatttctgctgaggacgatcggtccgccctctgggtgagtatctggctcggccttggcatcttcttggagaacgtagctgcacttgactgtgtggcgcggtatccaggacttttactttgaggaaattagagtgtttcaagcaggcgcacgccttgaatacattagcatggaataataagataggacctcggttctattttgttggtttctagagctgaggtaatgattaatagggatagttgggggcattcgtatttaattgtcagaggtgaaattcttggatttgttaaagacggactactgcgaaagcatttgccaaggatgttttca
>401b74102d854e9194b6e40deaf762a0788ba7f5_3
agctccaatagcgtatattaaagttgttgcggttaaaaagctcgtagttggatttctgccgaggacgaccggtccgcccactgggtgtgtatctggctcggcctgggcatcttcttggagaacgtagctgcacttgactgtgtggtgcggtatccaggacttttactttgaggaaattagggtgtttcaagcaggcgcacgccttgaatacattagcatggaataataagataggaccttggttctattttgttggtttctagaactgaggtaatgattaatagggatagttgggggcattcgtatttaactgtcagaggtgaaattcttggatttgttaaagacggactactgcgaaagcatttgccaaggatgttttca
>446be04449e7bd4dbb1332aab3d59b6a780bc37b_3
agctccaatagcgtatattaaagttgttgcggttaaaaagctagtagttggatttctgtcgaggacgaccggtccgccctccgggtgagtatctggttcggcctaggcatcctcttggagaaggagcggtcactttgctgtgaacgcccgtatccaggacttatactttgaggaaattggagtgcttcaagcaggcacacgccttgaacacatgagcatggaataatgcgataggacctcggttctattttgttggtttctagagctgaggtaatgataaatagggacagttgggggcattcgtatttaactgtcagaggtgaaattcttggatttgttaaagacggaccactgcgaaagcatttgccacggatgtcttca
>5a972385ad77370f1f3cd1533fafff808851a9c6_8
agctccaatagcgtatattaaagtggtggcggttaaaaagctcgtagttggatttctgccgaggacgaccggtccgcccactgggtgtgtatctggctcggcctgggcatcttcttggagaacgtatctgcacttgactgtgtggtgcggtatccaggacttttactttgaggaaattagagtgtttcaagcaggcacacgccttgaatacattagcatggaataataagataggaccttggttctattttgttggtttctagagctgaggtaatgattaatagggatagttgggggcattcgtatttaactgtcagaggtgaaattcttggatttgttaaagacggactactgcgaaagcatttgccaaggatgttttca
>60564a7afc8342d171cf4f8b8e96652b2e297768_3
agctccaatagcgtatattaaagttgttgcagttaaaaagctcgtagttgaatttctgctggagttgaataggtctgctctttgagtgtgtaccttatttcgacttcgcatcctcttagaatgttattaaagtactttattgtgctttatataatatctaagatctttactttgagtaaattatagtgtttcaagcaggcatattgccttgaatactttagcatggaataataaaataggattttggttctattttattggtttatagaactaaagtaatgattaatagggacagttgggggcattcgaatttaactgtcagaggtgaaattcttagatttgttaaagtcgaactactgcgaaagcatttgccaaggatgttttca
>6a7b045f7bddd28475c2d29a6c28e6a04573454a_3
cagctccaaaagcggatattaaagttgttgcagttaaaaagctcgtagttggattttatagtcgaagctacattttggtgtaactcgattatggtaataatttattattacacgttactttgaggaaattagagtgtttaaagcagatctttgtcatgtatatattagcatggaataacactaaggattattaaaactttgttggttactttttttaataatgattgatagggacagccgggggcatttgtatttaacagtcagaggtggaattcttggatttgttaaggacaaactattgcgaaagcatttgccaaggatgttttcg
>6f1a2a0b2723bb665470a0bc6b8e046171c1cc3b_3
agctccaatagcgtatattaaagttgttgcggttaaaaagctcgtagttggatttctgcttaggacgaccggtccgcccactgggtgagtatctggctcggcctgggcatcttcttggagaacgtagctgcgcttgattgtgtggtgcggaatccaggacttttactttgagaaaattagagtgtttcaagcaggcacacgccttgaatacattagcatggagtaataatataggacctcggttctattttgttggtttctagagctgaggtaatgattgatagggatagttgggggcattcgtatttaactgtcagaggtgaaattcttggatttgttaaagacggactactgcgaaagcatttgccaaggacgttccca
>70ccb08683d31f2554f0a6c1ebfe0f2ee3774d51_4
agctccaatagcgtatatttaagttgttgcagttaaaaagctcgtagttggatttctgccgaggacgaccggtccgcccactgggtgtgtatctggctcggcctgggcatcttcttggagaacgtagctgcacttgactgtgtggtgcggtatccaggacttttactttgaggaaattagagtgtttcaagcaggcacacgccttgaatacattagcatggaataataagataggaccttggttctattttgttggtttctagaactgaggtaatgattaatagggatagttgggggcattcgtatttaactgtcagaggtgaaattcttggatttttggaagacgaactactgcgaaagcatttaccaaggatgttttca
>7c5d0fa3a51c0c33ae346fc5fbf66eead60a292a_3
agctccaatagcgtatattaaagttgttgcggttaaaaagctcgtggttggatttctgctaaggacgaccggtccgccttcgggtgtgtacatgtgttgaccgaggcatcaatctggaagacgcctgtacttaaccgtgcgggcgtaattcagacgttttactttgaggaaattagagtgtttcaagcaggcaattgccctgaatacattagcatggaataataagataggacttcggttctattttgttggtttctagaactgaagtaataattaatagggacagtcgggggcattcgtactcaactgtcagaggtgaaattcttggatttgttgaagacgaactactgcgaaagcatttgccaaggatgttttca
>88512c3691291ff5821c4b2d945b6605ab9e6b05_4
agctccaggagcgtatattaaagttgttgcggttaaaaagctcgtagttggatttctgccaaggatgaccggtccgccttcgggtgtgtacctgtgttgtccgaggcatctttctggttggacgccagcacttcaatgtgttgtcgtactccagaacttttactttgaggaaattagagtgtttcaagcaggcatttgccttgaatacattagcatggaataataatctaggactccggttctattttgttggttctgagaacaggagtaatgattaatagggacagtcgggggcattcgtactcaactgtcagaggtgaaattcttggattagttggagacgaactactgcgaaagcatttgccaaggatgttttca
>954b122fd38f22d87765e04c226a8cf21cacca82_8
agctcgaagagcgtagattaaagttgttgcggttaaaaagctcgtagttggatttctgcttaggacgaccggtccgcccactgggtgagtatctggctcggcctgggcatcttcttggagaacgtagctgcgcttgattgtgtggtgcggaatccaggacttttactttgagaaaattagagtgtttcaagcaggcacacgccttgaatacattagcatggaataataatataggacctcggttctattttgttggtttctagagctgaggtaatgattgatagggatagttgggggcattcgtatttaactgtcagaggtgaaattcttggatttgttaaagacggactactgcgaaagcatttgccaaggatgttttca
>9b70b75a06a9230518b42e36244282ffce77cac0_3
agctccaatagcgtatattaaagttgttgcagttaaaaagctcgtagttgaatttctgctggagttgaataggtctgctctttgagtgtgtacctggtttcgacctggcatcctcttagaatgttattaaagtactttattgtgctttatataatatctaagatctttactttgagtaaattagagtgtttaaagcaggcatattgccttgaatactttagcatggaataataaaataggactttggttgtattttattggtttatagaactaaagtaatgattaatagggacagttgggggcattcgaatttaactgtcagaggtgaaattcttagatttgttaaagtcgaactactgcgaaagcatttgccaaggatgttttca
>b682ad855474e4a646f74f8cb5e4ec6ac2f5a7e0_3
agctccaatagcgtatatttaagttgttgcagttaaaaagctcgtagttggatttctgcttaggacgaccggtccgcccactgggtgagtatctggctcggcctgggcatcttcttggagaacgtagctgcgcttgattgtgtggggcggaatccaggacttttactttgagaaaattagagtgtttcaagcaggcacacgccttgaattcattagcatggaataataatataggacctcggttctattttgttggtttctagagctgaggtaatgattgatagggatagttgggggcattcgtatttaactgtcagaggtgaaattcttggatttgttaaagacggaatactgcgaaagcatttgccaaggatgttttca
>c1948c6861b48991d26efd8e8d36a3347cc302c0_14
agctctgatagtatatattaaagttgttgcagttaaaaagctcgtagttggatttctgctgaggacgatcggtccgccctctgggtgagtatctggctcggccttggcatcttcttggagaacgtagctgcacttgactgtgtggcgcggtatccaggacttttactttgaggaaattagagtgtttcaagcaggcgcacgccttgaatacattagcatggaataataagataggacctcggttctattttgttggtttctagagctgaggtaatgattaatagggatagttgggggcattcgtatttaattgtcagaggtgaaattcttggatttgttaaagacggactactgcgaaagcatttgccaaggatgttttca
>c29d5a6a583c4e4a62d2935b0e7f05c01fc19a27_3
agctccaatagcgtatattaaagttgttgcggttaaaaagctcgtagttagatttttgatctagtcgaccggtcactccaatggaatgtatcaggtttgattagaacatttgtctgaaatttttgtctgcacttcactgtgtggtcaaagattcagaccttttacattgaggaaattagagtgtttcaagcaggcatttgcattgaatacgttagcatggaataataaaataagactttggttttattttgttggtttctaaaactaaagtaatgattaatagggatagtcgggggcattcgtacttaactgtcagaggtgaaattcttggatttgttaaagacgaactactgcgaaagcatttgccaaggatgttttca
>c723baf8d9e69bfcb43ec4c460e8b09f370c74a2_5
agctccaatagcgtatattaaagttgttgcagttaaaaagctcgtagttgaatttctgctggagttgaataggtctgctcttagagtgtgtacattgtttcgacttcgcatcctcttagaatgttattaaagtactttattgtgctttatataatatctaagatctttactttgagtaaattagagtgtttcaagcaggcatatcgccttgaatactttagcatggaataataaaataggattttggttctattttattggtttatagaactaaagtaatgattaatagggacagttgggggcattcgaatttaactgtcagaggtgaaattcttagatttgttaaagtcgaactactgcgaaagcatttgcaaaggatgttttca
>cda199453e3ec8a2d64ba763aba8796c2c0e5d71_3
agctccaatagcgtatattaaagttgttgcagttaaaaagctcgtagttgaatttctgctggagttgaataggtctgctctttgagtgtgtaccttgtttcgacttcgcatcctcttagaatgttattaaagtactttattgtgctttatataatatctaagatctttactttgagaaaattagagtgtttcaagcaggcatattgccttgaatactttagcatggaataataaaataggattttggttctattttattggtttatagaactaaagtaatgattaatagggacagttgggggcattcgaatttaactgtcagaggtgaaattcttagatttgttaaagtcgaactactgcgaaagcatctgccaagaatgttccca
>cf3a4dffe7562616af133ac9fd059bcd4139d5cc_3
agctccaaaagcatatattaaagttgttgcggttaaaaagctcgtagttggattttatagtcggagctacattttggtgtaactcgattatggtaataatttattattacacgttactttgaggaaattagagtgtttaaagcagatctttgtcatgtatatattagcatggaataacactaaggattattaaaactttgttggttactttttttaataatgattgatagggacagccgggggcatttgtatttaacagtcagaggtgggattcttggatttgttaaggacaaactattgcgaaagcatttgccaaggatgttttcg
>cf71e0a1e86f46009c46d8805d2f6b63c84f4d74_4
agctccaatagcgtatatttaagttgttgcagttaaaaagctcgtagttggatttctgccgaggacgaccggtccgccctctgggtgagtatctggctcggcctgggcatcttcttggagaacgtatctgcacttgactgtgtggtgcggtattcaggacttttactttgaggaaattagagtgtttcaagcaggcatacgccttgaatacattagcatggaataataagataggacctcggttctattttgttggtttctagagctgaggtaatgattgatagggatagttgggggcattcgtatttaactgtcagaggtgaaattcttggatttgttaaagacggactactgcgaaagcatttgccaaggatgttttca
>d3a4908bfff595702defa00d9dbc8e5eed1dd52a_3
agctccaatagcgtatattaaagttgttgcagttaaaaagctcgtagttgaatttctgctggagttgaataggtctgctctttgagtgtgtaccttgtttcgacttcgcatcctcttagaatgttattaaagtactttattgtgctttatataatatctaagatctttactttgagaaaattagagtgtttcaagcaggcatattgccttgaatactttagcatggaataataaaataggattttggttctattttattggtttatagaactaaagtaatgattaatagggacagttgggggcattcgaatttaactgtcagaggtgaaattcttagatttgcggaagactaactagagcgaaagcatttgccaaggatgttttca
>ece7cdd1092f082ce4f453691ce958c73d387253_3
agctccaaaagtgtatattaaagttgttgcggttaaaaagctcgcagttggattttatagtcggagctacattttggtgtaactcgattatggtaataatttattattacacgttactttgaggaaattagagtgtttaaagcagatccttgtcatgtatatattagcatggaataacactaaggattgttaaaactttgttggttactttttttaataatgattgatagggacagccgggggcatttgtatttaacagtcagaggtggaattcttggatttgttaaggacaaactattgcgaaagcatttgccaaggatgttttcg
>f023571e62541334f53cd871436d376e43822c04_5
agctccaatagcgtatattaaagttgttgcggttaaaaagctcgtagttggatttctgctgaggacgaccggtccgccctctgggtgagtatctggctcggccttggcatcttcttggagaacgtagctgcacttgactgtgtggtgcggtatccaggacttttactttgaggaaattagagtgtttcaagcaggcacacgccttgaatacattagcatggaataataagataggaccttggttctattttgttggtttctagaactgaggtaatgattaatagggatagttgggggcattcgtatttaactgtcagaggtgaaattcttggatttgttaaagacggactactgcgaaagcatttgccaaggatgttttca
>f6e387974264f02dd3fc45e7658376090ef0e0f1_5
agctctgatagtatatattaaagttgttgcggttaaaaagctcgtagttggatttctgctgaggacgaccggtccgccctctgggtgagtatctggctcggccttggcatcttcttggagaacgttactgcacttgattgtgtggtgcggtatccaggacttttactttgaggaaattagagtgtttcaagcaggcgcacgccttgaatacattagcatggaataataagataggaccttggttctattttgttggtttctagagctgaggtaatgattaatagggatagttgggggcattcgtatttaactgtcagaggtgaaattcttggatttgttaaagacggactactgcgaaagcatttgccaaggatgttttca
>0c94ca471ba42c6ed756c6d95bdcce6f8a6a665a_4
agctctgatagtatatattaaagttgttgcagttaaaaagctcgtagttgaatttctgctggagttgaataggtctgctctttgagtgtgtaccttgtttcgacttcgcatcctcttagaatgttattaaagtactttattgtgctttatataatatctaagatttttactttgagtaaattagagtgtttaaagcaggcatattgccttgaatactttagcatggaataataaaataggactttggttctattttattggtttatagaactaaagtaatgattaatagggacagttgggggcattcgaatttaactgtcagaggtgaaattcttagatttgttaaagtcgaactactgcgaaagcatttgccaaggatgttttca
>12adbe05d11abbf6fe5db52cc8c215c39238898b_4
agctccaatagcgtatattaaagttgttgcggttaaaaagctcgtagttggatttctgccgaggacgaccggtccgcccactgggtgtgtatctggctcggcctgggcatcttcttggagaacgtagctgcacttgactgtgtggtgcggtatccaggacttttactttgaggaaattagagtgtttcaagcaggcacacgccttgaatacattagcatggaataataagataggaccttggttctattttgttggtttctagaactgaggtaatgattaatagggatagttgggggcattcgtatttaattgtcagaggtgaaattcttggatttgttaaagacggactactgcgaaagcatttaccaaggatgttttca
>184630ce3169cedb863a2b41146a76d4184a382d_6
agctccaatagcgtacattaaagttgttgcggttaacaagctcgtagttggatttctgccgaggacgaccggtcggccctctgggtttgtctctggctcggcctgggcatcttcttggagaccgtaaccgcacttgactgtgtggtgcggtattcaggacttttactttgaggaaattggagtgtttcaagcaggcacacgccttgaatacattagcatggaaaagaagataggacctcggttctattctgttggcttctagagctgaggtaatgattgatagggatagttgggggcattcgtatttaactgtcagaggtgaaattcttggatttgttaaagacggactactgcgaaagcatttgccaaggatgttttca
>1b0fca2238f52d44bd4512120464f6b31fcfa462_3
agctccaatagcgtatattaaaattgttgcagttaaaaagctcgtagttgaatttctgctggagttgaataggtctgctctttgagtgtgtaccttgtttcgacttcgcatccccttagaatgttattaaagtactttattgtgctttatataatatctaagatttttactttgagtaaattagagtgtttaaagcaggcatattgccctgaatactttagcatggaataataaaataggactttggttctattttattggtttatagaactaaagtaatgattaatagggacagttgggggcattcgaatttaactgtcagaggtgaaattcttagatttgttaaagtcgaactactgcgaaagcatttgccaaggatgttttca
>1d9f8af9ee85e3ca41d87fff16ba2b18a86c99ab_3
agctctgatagtatatattaaagttgttgcggttaaaaagctcgtagttggatttctgctgaggacgaccggtccgccctctgggtgagtatctggctcggccttggcatcttcttggagaacgttactgcacttgattgtgtggtgcggtatccaggacttttactttgaggaaattagagtgtttcaagcaggcgcacgccttgaatacattagcatggaataatagaataggacgtcgtttctattttgttggttttcggaaatcgacgtaatgattaatagggatagttgggggcattcgtatttaattgtcagaggtgaaattcttggatttgttaaagacggactactgcgaaagcatttgccaaggatgttttca
>230e6666cf24a7144b9cfd89607c2ca78bef5879_3
atctccaaaagcgtatattaaagttgttgcggttaaaaagctcgcagttggattttatagtcggagctacattttggtgtaactcgattatggtaataatttattattacacgttactttgaggaaattagagtgtttaaagcagatctttgtcatgtatatattagcatggaataacactaaggattattaaaactttgttggttactttttttaataatgattgatagggacagccgggggcatttgtatttaacagtcagaggtggaatccttggattcgttaaggacaaactattgcgaaagcatttgccaaggatgttttcg
>29b4f63d2d73a98a16e5c91d8bffed5848101a15_3
agctccaatagcgtatattaaagttgttgcagttaaaaagctcgtagttgaatttctgctggagttgagtaggtctgctctttgagtgtgtaccttgtttcgacttcgcatcctcttagaatgttattaaagtactttattgtgctttatataatatctaagatttttactttgagtaaattagagtgtttcaagcaggcatattgccttgaatactttagcatggaataataagatgggattttggttctattttattggtttatagaactaaagtaatgattaatagggacagttgggggcattcgaatttaactgtcagaggtgaaattcttagatttgttaaagtcgaactactgcgaaagcatttgccaaggatgctttca
>32657cf7f28fc0772b6f408eeeca3a3d0938ac7e_3
agctccaatagcgtatcttaatgttgttgcagttaaaaagctcgtagttggatttctgccgaggacgaccggtccgcccactgggtgtgtatctggctcggcctgggcatcttcttggagaacgtagctgcacttgactgtgtggtgcggtatccaggacttttactttgaggaaattagagtgtttcaagcaggcacacgccttgaatacattagcatggaataataagataggaccttggttctattttgttggtttctagaactgaggtaatgattaatagggatagttgggggcattcgtatttaactgtcagaggtgaaattcttggatttgttaaagacggactactgcgaaagcatttgccaaggatgttttca
>3f5916dbedace7fe186b6695c3853f5d611ff4b5_4
agctccaatagcgtatattaaagttgttgcggttaaaaagctcgtagttggatttctgcttaggacgaccggtccgcccactgggtgagtatctggctcggcctgggcatcttcttggagaacgtagctgcgcttgattgtgtggtgcggaatccaggacttttactttgagaaacttagagtgtttcaagcaggcacacgccttgaatacattagcatggaataataatataggacctcggttctattttgttggtttccagagctgaggtaatgattgatagggatagttgggggcattcgtatttaactgtcagaggtgaaattcttggatttttggaagacggactactgcgaaagcatttgccaaggatgttttca
>40d384cca0e8c2905e7b9c6100268cb4404a9be2_3
agctccaatagcgtatattaaagttgttgcagttaaaaagctcgtagttgaatttctgcgagagttgaataggtctgctctttgagtgtgtaccttatttcgacttcgcatcctcttagaatgctataaaagtactttattgtgctttttataaaatctaagatttttactttgagtaaattagagtgtttcaagcaggcatattgccttgaatactttagcatggaataataagataggactttggttctattttgttggtttatagaactaaagtaatgattaatagggacggttgggggcattcgaatttaactgtcagaggtgaaattcttagatttgttaaagtcgaactactgcgagagcatttgccaaggatgttttca
>455fa1e124a3f88550a9bc01f0a634643a298094_3
agctccaatagcgtatattaaagttgttgcagttaaaaagctcgtagttgaatttctgctggagttgattaggtctgctctgtgagtgtgtaccttaatatcgacttagcatcctcttagaatgttattaaagtactttattgtgctttatataatatctaagatttttactttgagtaaattagagtgtttatagcaggcatattaccttgaatactttagcatggaataataaaataggattttggttctattttattggtttatagaactaaagtaatgattaatggggacagttgggggcattcgaatttaactgtcagaggtgaaattcttagatttgttaaagtcgaactactgcgaaagcatttgccaaggatgttttca
>48896daee6d0282204948ba4f1eccecfd778d12e_3
agctccaatagcgtatattaaagttgttgcggttaaaaagctcgtagttggatttctgccgaggacgaccggtccgcccactgggtgtgtatctggctcggcctgggcatcttcttggagaacgtatctgcacttgactgtgtggtgcggtatccagaacttttactttgaggaaattagagtgtttcaagcaggcacacgccttgaatacattagcatggagtaataagataggaccttggttctattttgttggtttctagagctgaggtaatgattaatagggatagttgggggcattcgtatttaactgtcagaggtgaaattcttggatttgttaaagacggactactgcgaaagcatttgccaaggatgttttca
>4b9a7f37eebfa2bb39df44615dd57fa3013d7e9e_4
agctccaatagcgtatattaaagttgttgcagttaaaaagctcgtagttgaatttctgctggagttgaatagttcggctctttgagtgtgtaccttgtttcgacttcgcatcctcttagaatgttattaaagtactttattgtgctttatataatatctaagatttttactttgagtaaattagagtgtttcaagcaggcatattgccttgaatactttagcatggaataataagataggattttggttctattttattggtttatagaactaaagtaatgattaatagggacagttgggggcattcgaatttaactgtcagaggtgaaattcttagatttgttaaagtcgaactactgcgaaagcatttgccaaggatgttttca
>555654bcf76e644fd57214a89ff851446edee952_5
agctccaatagcgtatattaaagttgttgcggttaaaaagctcgtagttggatttctgccgaggacgaccggtccgccctctgggtgagtatctggctcggcctgggcatcttcttggagaacgtagctgcgcttgattgtgtggggcggaatccaggacttttactttgagaaaattagagtgtttcaagcaggcacacgccttgaatacattagcatggaataataatataggacctcggttctattttgttggtttctagagctgaggtaatgattgatagggatagttgggggcattcgtatttaactgtcagaggtgaaattcttggatttgttaaagacggactactgcgaaagcatttgccaaggatgttttca
>73542d022ad821bebf391c891e25bb9126bc20d9_3
agctctgatagtatatattaaagttgttgcagttaaaaagctcgtagttggatttctgcttaggacgaccggtccgcccactgggtgagtatctggctcagccttggcatcttcttggagaacgtagctgcgcttgattgtgtggtgcggaatccaggacttttactttgagaaaattagagtgtttcaagcaggcacacgccttgaatacattagcatggaataataatataggacctcggttctattttgttggtttctagagctgaggtaatgattgatagggatagttgggggcattcgtatttaactgtcatagttgaaattcttggatttgttaaagacggactactgcgaaagcatttgccaaggatgttttca
>73daf1f9b09b68471ccc8848c3ad7ccede1b9001_4
agctccaatagcgtatattaaagttgttgcggttaaaaagctcgtagttggatttctgcttaggacgaccggtccgcccactgggtgagtatctggctcggccttggcatcttcttggagaacgttactgcacttgattgtgtggtgcggtatccaggacttttactttgaggaaattagagtgtttcaagcaggcacacgccttgaatacattagcatggaataataagataggaccttggttctattttgttggtttctagaactgaggtaatgattaatagggatagttgggggcattcgtatttaactgtcagaggtgaaattcttggatttgttaaagacggactactgcgaaagcatttgccaaggatgttttca
>77208a3acdf9229dc60582d4adfad6401d0e99a3_5
agctccaaaagcgtatattaaagttgttgcggttaaaaagctcgtagttggattttatagtcgaagctacattttggtgtaactcgattatggtaataatttattattacacgttactttgaggaaattagagtgtttaaagcagatctttgtcatgtatatattagcatggaataacactaaggattattaaaactttgttggttactttttttaataatgattgatagggacagccgggggcatttgtatttaacagtcagaggtggaattcttggatttgttaaggacaaactattgcgaaagcatctgccaagaatgttccca
>78c59b6a3242068436465adb5fa041ea638fcae2_5
agctccaatagcgtatattaaagttgttgcagttaaaaagctcgtagttggatttctgctgaggacgaccggtccgccctctgggtgagtatctggctcggccttggcatcttcttggagaacgttactgcacttgattgtgtggtgcggtatccaggacttttactttgaaaaaattagagtgtttcaagcaggcgttttgctatgaatacattagcatggaataataagataggacctcggttctattttgttggtttctagagctgaggtaatgattaatagggatagttgggggcattcgtatttaactgtcagaggtgaaattcttggatttgttaaagacggactactgcgaaagcatttgccaaggatgttttca
>7b535fbe7351a10f07fa5aa08f0e7f70fd50a7de_3
agctccaatagcgtatattaaagttgttgcagttaaaaagctcgtagttggatttctgcttaggacgaccggtccgcccactgggtgagtatctggctcggcctgggcatcttcttggagaacgtagctgcgcttgattgtgtggggcggaatccaggacttttactttgagaaaattagagtgtttcaagcaggcacacgccttgaatacattagcatggaataataatataggacctcggttctattttgttggtttctagagctgaggtaatgattgatagggatagttgggggcattcgtatttcatagtcagaggtgaaattcttggatttgttaaagacggactactgcgaaagcatttgccaaggatgttttca
>84a36f1337864c5bef84d903d8d4dbfd25cfad1d_3
agctccaataggatatattaaaattgttgcagttaaaaagctcgtagttgaatttctgctggagttgaataggtctgctctttgagtgtgtaccttgtttcgacttcgcatcctcttagaatgttattaaagtactttattgtgctttatataatatctaagatttttactttgagtaaattagagtgtttaaagcaggcatattgccttgaatactttagcatggaataataaaataggactttggttctattttattggtttatagaactaaagtaatgattaatagggacagttgggggcattcgaatttaactgtcagaggtgaaattcttagatttgttaaagtcgaactactgcgaaagcatttgccaaggatgttttca
>8816255c36035eb9c73eceb33b50e546a27f46cd_4
agctccaatagcgtatattaaagttgttgcggttaaaaagctcgtagttggatttctgcttaggacgaccggtccgcccactgggtgagtatctggctcggcctgggcatcttcttggagaacgtatctgcacttgactgtgtggtgcggtattcaggacttttactttgaggaaattagagtgtttcaagcaggcatacgccttgaatacattagcatggaataataagataggacctcggttctattttgttggtttctagatctgaggtaatggttaatagggatagttgggggcattcgtatttaactgtcagaggtgaaattcttggatttgttaaagacggactactgcgaaagcatttgccaaggatgttttca
>8a7145b6d0d688f4e925c91ea792df5898efd471_3
agctccaatagcgtatattaaagttgctgcggttaaaaagctcgtagttggatttctgctgaggaagaccggtctgccctctgggtgagtatctggctccgccttggcatcttcttgggaaccgttgctgcacttgattgtgtggtgcgggattcaggacatttactttgaggaaattagagtgcttcaagaaggcatggtgtctgaatacattagcatggaataataagatgggccctggctctgttttgttggtttctggagccgaggtcatgattaacagggacaattgggggcattcgtacttaactgtcagaggtgaaattcttggatttgttaatgacggactactgcgaaagcatttgccaaggatgttttca
>909206944c59a4e31b48413b9a11c5c39a206bd2_3
agctctgatagtatatattaaagttgttgcggttaaaaagctcgtagttggatttctgttgaggacgaccggtccgccctctgggtgagtatctggctcggccttggcatcttcttggggaacgttactgcacttgactgtgtggtgcggtatccaggacttttactttgaggaaattagagtgtttcaagcaggcgcacgccttgaatacattagcatggaataataagataggacctcggttctattttgttggtttctagagctgaggtaatgattaatagggatagttgggggcattcgtatttaactgtcagaggtgaaattcttggatttgttaaagacggactactgcgaaagcatttgccaaggatgttttca
>941e7fb35a1187036e96695bb81d98f64f91fbbf_4
agctccaatagcgtatattaaagttgttgcggtcaaaaagctcgtagttggatttctgccgaggacgaccggtccgcccactgggtgtgtatctggcccggcctgggcatcttcttggagaacgtagctgcacttgactgtgcggtgcggtatccaggacttttactttgaggaaattagagtgtttcaagcaggcacacgccttgaatacattagcatggaataataagataggaccttggttctattttgttggtttctagagctgaggtaatgattgatagggatagttgggggcattcgtatttaactgtcagaggtgaaattcttggatttgttaaagacggactactgcgaaagcatttgccaaggatgttttca
>a3bf44780ae346ada7cd1e490c47e606a7127abf_4
agctccaatagcgtatattaaagttgttgcggttaaaaagctcgtagttggatttctgctgaggacgatcggtccgccctctgggtgagtatctggctcggccttggcatcttcttggagaacgtagctgcacttgactgtgtggcgcggtatccaggacttttactttgagaaaattagagtgtttcaagcaggcacacgccttgaatacattagcatggaataataatataggacctcggttctattttgttggtttctagagctgaggtaatgattgatagggatagttgggggcattcgtatttaactgtcagaggtgaaattcttggatttgttaaagacggactactgcgaaagcatatgccaaggatgttttca
>b1093edf2c320761928a2656331c1cd92815e6e2_4
agctccaatagcgtatattaaagttgttgcggttaaaaagctcgtagttggatttctgcttaggacgaccggtccgccctctgggtgagtatctggctcggccttggcatcttcttggggaacgttactgcacttgactgtgtggtgcggtatccaggacttttactttgaggaaattagagtgtttcaagcaggcgcacgccttgaatacattagcatggaataataagataggacctcggttctattttgttggtttctagagctgaggtaatgattaatagggatagttgggggcattcgtatttaactgtcagaggtgaaattcttggatttgttaaagacggactactgcgaaagcatttgccaaggatgttttca
>bfda99444b9daa2d2ab0ede9bda49c5da1ff96d2_3
agctccaatagcgtatattaaatttgttgcggttaaaaagctcgtagttggatttctgccgaggacgaccggtccgccctctgggtgagtatctggctcggccttggcatcttcttggggaacgttactgcacttgactgtgtggtgcggtatccaggacttttactttgaggaaattagagtgtttcaagcaggcgcacgccttgaatacattagcatggaataataagataggacctcggttctattttgttggtttctagagctgaggtaatgattaatagggatagttgggggcattcgtatttaactgtcagaggtgaaattcttggatttgttaaagacggactactgcgaaagcatttgccaaggatgttttca
>c311ef5d8908092d7649a4b86ff34273105b6a6e_3
agctccaaaagcgtatattaaagttgttgcggttaaaaagctcgtagttggattttatagtcgaagcaacattttggtgtaactcgattatggtaataatttattattacacgttactttgaggaaattagagtgtttaaagcagatctttgtcatgtatatattagcatggaataacactaaggattattaaaactttgttggttactttttaataatgattgatagggacagccgggggcatttgtatttaacagtcagagatggaattcttggatttgttaaggacaaactattgcgaaagcatttgccaaggatgttttcg
>cb0e88950a2cc289413931c33489bdbbb2df962c_3
agctccaatagcgtatattaaagttgttgcggttaaaaagctcgtagttggatttctgcttaggacgaccggtccgcccactgggtgagtatctggctcggcctgggcatcttcttggagaacgtagctgcgcttgattgtgtggtgcggaatccaggacttttactttgagaaaattagagtgtttcaagcaggcacacgccttgaatacattagcatggagtaataatataggacctcggttctattttgttggtttctagagctgaggtaatgattgatagggatagttgggggcattcgtatttcatagtcagaggtgaaattcttggatttgttaaagacggactactgcgaaagcatttgccaaggatgttttca
>ce9456ca3f4ce42c2991fb9de99546e195028d77_3
agctccaatagcgtatattaaatttgttgcggttaaaaagctcgtagttggatttctgccgaggacgaccggtccgcccactgggtgtgtatctggctcggcctgggcatcttcttggagaacgttactgcacttgactgtgtggtgcggtatccaggacttttactttgaggaaattagagtgtttcaagcaggcgcacgccttgaatacattagcatggaataataagataggacctcggttctattttgttggtttctagagctgaggtaatgattaatagggatagttgggggcattcgtatttaactgtcagaggtgaaattcttggatttgttaaagacggactactgcgaaagcatttgccaaggatgttttca
>e96580a9c02b0178f975b96fdee24b6652502d0b_4
agctccaatagcgtatattaaagttgttgcagttaaaaagctcgtagttgaatttctgctggagttgaataggtctgctctttgagtgtgtaccttgtttcgacttcgcatcctcttagaatgttattaaagtactttattgtgctttatatagtatctaagatctttactttgagaaaattagagtgtttcaagcaggcatattgccttgaatacattagcatggaataataaaataggattttggttctattttattggtttatagaactaaagtaatgattaatagggacagttgggggcattcgaatttaactgtcagaggtgaaattcttagatttgttaaagtcgaactactgcgaaagcatttgccaaggatgttttca
>ebdeec8e1685a0f5f47bf0781aab60d9dd0247a4_7
agctccaatagcatatattaaagttgttgcggttaaaaagctcgtagttggatttctgctgcgaatgaccggtccgccttccgggtgagtatctggttcgttcgtggcattttctcggagaatgtagctgcacttgactgtgtggtgcgccatctgggacttttactttgaggaacttagagtgcttcaagcaggccagtgccttgaatacattagcatggaataataagataggaccttggttttattttgctggtttctaaagctgaggtaacgattgatagggacagttgggggcattcgtacttaactgtcagaggtgaaattcttggatttgttaaagacggaccgctgcgaaagcatttgccaaagatgttttca
>fada23666e1108b91be2e58113946ded4de67a67_3
agctccaatagcgtatattaaagttgttgcagttaaaaagctcgtagttggatttctgcttaggacgaccggtccgcccactgggtgagtatctggctcggcctgggcatcttcttggagaacgtagctgcgcttgattgtgtggggcggaatccaggacttttactttgagaaaattagagtgtttcaagcaggcacacgccttgaatacattagcatggaataataatataggacctcggttctattttgttggtttctagagctgaggtaatgattgatagggatagttgggggcattcgtatttaactgtcagaggtgaaattattggatttgttaaagacgaacaactgcgaaagcatttgccaaggatgttttca
>07addf23dc12ac3650d6ac9a08d5a601263a2ea2_3
agctccaggagcgtatattaaagttgttgcagttaaaaagctcgtagttggatttctgctgaggcctaacggtcagcccattgggtttgcatctgctacggcctgagcatcttcttggagaaggttgctgcacttgactgtgtggcgcggtatccaggacttttactttgaggaaattagagtgcttcaagcaggcacacgccttgaacacgtgagcatggaataatgtgataggacctcggttctattttgttggtttctagagctgaggtaatgataaatagggacagttgggggcattcatatttaactgtcagaggtgaaattcttggatttgttaaagatggactactgcgaaagcatttgccacggatgtcttca
>10e973bc2abbde367ebe10474e15c9612f44bc1f_5
agctccaatagcgtatattaaagttgttgcggttaaaaagctcgtagttggatttctgcttaggacgaccggtccgcccactgggtgagtatctggctcggcctgggcatcttcttggagaacgtagctgcgcttgattgtgtggtgcggaatccaggacttttactttgagaaaattagagtgtttcaagcaggcacacgccttgaatacattagcatggaataataatataggacctcggttctattttgttggtttctagagctgaggtaatgattgatagggatagttgggggcattcgtatttaactgtcagaggtgaaattcttggatcagttgaagactaactactgcgaaagcatttgccaaggatgttttca
>174b1e886c97053dd51e51c610abead552c834ef_4
agctccaatagcgtatattaaagttgttgcagttaaaaagctcgtagttgaatttctgctggagttgaataggtctgctctttgagtgtgtaccttgtctcgacttcgcatcctcttagaatgttattaaagtactttattgtgctttatataatatctaagatttttactttgagtaaattagagtgtttaaagcaggcatattgccttgaatactttagcatggaataataaaataggactttagttctattttattggtttatagaactaaagtaatgattaatagggacaattgggggcattcgaatttaactgtcagaggtgaaattcttagatttgttaaagtcgaactactgcgaaagcatttgccaaggatgttttca
>17cfad4daaea8cd9953a25698b664323831d3ce2_3
agctccaatagcgtatattaaagttgttgcagttaaaacgctcgtagttgaatttctgctggagttgaataggtcggctctttgagtgtgtaccttgtttcgacttcgcatcctcttagaatgttattaaagtactttattgtgctttatataatatctaagatttttactttgagtaaattagagtgtttaaagcaggcatattgccttgaatactttagcatggaataataaaataggactttagttctattttattggtttatagaactaaagtaatgattaatagggacagttgggggcattcgaatttaactgtcagaggtgaaattcttagatttgttaaagtcgaactactgcgaaagcatttgccaaggatgttttca
>1f1190ecbb7bb604bcee174d8c547bced08c564a_5
agctctgatagtatatattaaagttgttgcggttaaaaagctcgtagttggatttctgccgaggacgaccggtccgccctctgggtgcgtatctggctcggcctgggcatcttcttggagaacgtatctgcacttgactgtgtggtgcggtatccaggacttttactttgaggaaattagagtgtttcaagcaggcatacgccttgaatacattagcatggaataataagataggacctcggttctattttgttggtttctagagctgaggtaatgattaatagggatagttgggggcattcgtatttaactgtcagaggtgaaattcttggatttgttaaagacggactactgcgaaagcatttgccaaggatgttttca
>2cd9a5773ee03a3cfc94f7370bc8cf5470ef53b7_1
agctccaatagcgtatattaaagttgttgcggttaaaaagctcgtagttggatttctgctgaggacgaccggtccgccctctgggtgcgtatctggctcggccttggcatcttcttggagaacgttactgcacttgactgtgtggtgcggtatccaggacttttactttgaggaaattagagtgtttcaagcaggcatacgccttgaatacattagcatggaataataagataggaccttggttctattttgttggtttctagagctgaggtaatgattaatagggatagttgggggcattcgtatttaactgtcagaggtgaaattcttggatttgttaaagacggactactgcgaaagcatttgccaaggatgttttca
>3132bf53ed7c7646cdd8501672cf38ac4590eeae_4
agctctgatagtatatattaaagttgttgcagttaaaaagctcgtagttgaatttctgctggagttgaataggtctgctctttgagtgtgtaccttgtttcgacttcgcatcctcttagaatgttattaaagtactttattgtgctttatataataactaagatctttactttgagaaaattagagtgtttcaagcaggcatattgccttgaatactttagcatggaataataaaataggattttggttctattttattggtttatagaactaaagtaatgattaatagggacagttgggggcattcgaatttaactgtcagaggtgaaattcttagatttgttaaagtcgaactactgcgaaagcatttgccaaggatgttttca
>3a9ee8362853e6d4bf9798ad6dc28e740bd35618_3
agctccaatagcgtatatttaagttgttgcagttaaaaagctcgtagttggatttctgcttaggacgaccggtccgcccttgagcatctggctcggcctgggcatcttcttggagaacgtagctgcgcttgattgtgtggtgcggaatccaggacttttactttgagaaaattagagtgtttcaagcaggcacacgccttgaatacattagcatggaataataatataggacctcggttctattttgttggtttctagagctgaggtaatgattgatagggatagttgggggcattcgtatttaactgtcagaggtgaaattcttggatttactgaagactaactactgcgaaagcatttgccaaggatgttttca
>4a2def16f7ec2c2acd8c9a6d338695cc8622f892_3
agctccaggagcgtatattaaagttgttgcagttaaaacgctcgtagttggatttctgctgaggacgaccggtccgccctctgggtgagcatctggctcggcctgagcatcctcttggagaaggtgtgtgcactttactgtgtgcacccgtatccaggacttttactttgaggaaattggagtgcttcaagcaggcacacgccttgaacacatgagcatggaataatgcgataggacctcggttctattttgttggtttctagaactgaggtaatgataaatagggacagttgggggcattcgtatttaactgtcagaggtgaaattcttggatttgttaaagacggaccactgcgaaagcatttgccacggatgtcttta
>4e9180a82d2b0c7419c23600e56d28079de5c9ba_5
agctccaatagggtattttaaaggtggtgcggttaaaaagctcgtagttggatttctgcttaggacgaccggtccgcccactgggtgagtatctggctcggcctgggcatcttcttggagaacgtagctgcgcttgattgtgtggtgcggaatccaggacttttactttgagaaaattagagtgtttcaagcaggcacacgccttgaatacattagcatggaataataatataggacctcggttctattttgttggtttctagagctgaggtaatgattgatagggatagttgggggcattcgtatttaactgtcagaggtgaaattcttggatttgttaaagacggactactgcgaaagcatttgccaaggatgttttca
>94bc4dff1b587b35c04e74d3f0f65453509fac59_1
agctccaatagcgtatattaaagttgttgcggttaaaaagctcgtagttggatttctgccgaggacgaccggtccgccctctgggtgcgtatctggctcggcctgggcatcttcttggagaacgtatctgcacttgactgtgtggtgcggtatccaggacttttactttgaggaaattagagtgtttcaagcaggcgcacgccttgaatacattagcatggaataataagataggacctcggttctattttgttggtttctagagctgaggtaatgattaatagggatagttgggggcattcgtatttaattgtcagaggtgaaattcttggatttgttaaagacggactactgcgaaagcatttgccaaggatgttttca
>97ad310530563be46378d9f479194c27c6ce10a1_1
agctccaatagcgtatattaaagttgttgcggttaaaaagctcgtagttggatttctgccgaggacgaccggtccgccctccgggtgagtatctggctcggcctgggcatcttcttggagaacgtagctgcacttgactgtgtggtgcggtatccaggacttttactttgaggaaattagagtgtttcaagcaggcacacgccttgaatacattagcatggaataataagataggacctcggttctattttgttggtttctagagctgaggtaatgattaatagggatagttgggggcattcgtatttaactgtcagaggtgaaattcttggatttgttaaagacggactactgcgaaagcatttgccaaggatgttttca
>d23d7a20f58396dcc5c4bf2f5e0fff05c9248d8a_3
agctccaatagcgtatattaaagttgttgcggttaaaaagctcgtagttagatttttgacctagtcgaccggtcactccaatggaatgtatcaggtttgattaggttatttgtctgatatttttggctgcacttgactatgtggtcaaatgttcagaccttttactttgaggaaattagagtgtttcaagcaggcatttgccttgaatacattagcatggaataataaaataagactttggttttattttgttggtttctaaaactaaagtaatgattaatagggatagtcgggggcattcgtacttaactgtcagaggtgaaattcttggatttgttaaagacgaactactgcgaaagcatttgccaaggatgttttca
>d42a0ed9e5df2219bdbe74578afc678e70005539_3
ggctctagtagcgtatattaaagttgttgcagttaaaaagctcgtagttgaatttctgctggagttgaataggtctgctctttgagtgtgtaacttgtttcgacttcgcatcctcttagaatgttattaaagtactttattgtgctttatataatatctaagatttttactttgagtaaattagagtgtttcaagcaggcatattgccttgaatactttagcatggaataataaaataggattttggttctattttattggtttatagaactaaagtaatgattaatagggacagttgggggcattcgaatttaactgtcagaggtgaaattcttagatttgttaaagtcgaactactgcgaaagcatttgccaaggatgttttca
