## Supplementary material for "The windblown: possible explanations for dinophyte DNA in forest soils": File S2

d6a793bd001362ed30a1ba5f8a06fa8dd60561c7_19311
b8e2af4b0ca8de85d4643140f299e0a6a5eb1e51_17104
4c554190d2d6dd436de4e6c438462f619e18265f_5365
fc96e6a67377f73fe72f4d4638fbec6c59194dc3_4520
7ee5cf32c634bd73780636ced48ccf80b5484854_4826
f9e3bb9f1cc7d1d39480a29bb220c52bc9b27608_2933
93a188b7435e9ac5053755f2989f2cca340dba24_3263
402ff1c02cf1114cb6b6cfe542997b9d276dedbd_2668
92dac336af3373c32854d83a7535c7232bb2e264_2893
e26590a7c401a8e17ddf3090947520f9f3c1fea3_2967
d1f1e1ed2c659894415f7e0408f5dae1d6ecad2c_1593
f5390614fe3126281636dd5b8fa4cffaf8c3b26c_1991
f5d50cc840e1144e93eeaa6d29b067d9ac79902e_1764
720928ac158e8a47cbdb1c68d0bb65aed01439e5_1788
793ed5f0b15e46106a691ff2ff3bb4ca11033d3e_2838
5e197728f5006b8e36139eabb3d4562d4c5df937_1488
6698fbde9ac45aba6fab523c102479eb1fa4988f_953
4d66b77aead4818a5d3d13f9f27ee17d2164f1d7_987
aba964e993016c42c28a49f572a9176a7424e41e_913
4927cf57794970028dd04aa6cbd89a4937834072_566
9d9b7584bef62b3c85d708da59bb2f6556d5efde_505
1ce825f5402141d6410f1766f3916e5c6d943c59_840
a0fd5c3584dad6151533e2e5545ecd105c622188_388
e1e8bdf388cb28c352b31143811a618cee79ef77_373
0e13994ab91bce37f31dad5fd0949ed600a4b434_360
c26186e453d2655a42433bc36cf20b587a21a457_526
856a5e57aeaf7fbc8e554ef054a7f8d1dc1c7419_587
8deba22c1e1d6883a2b33a0bf42c989cbd5fca23_643
78d3f5bf16caf870fd8b2510c52ae4d38b48e8fe_359
98a6d25c852f97c65fca4e09121b3f88756272f9_310
ee99783001b9b4c561ea052f5355a01588c49ddb_207
6c9c00178e26af4f55cdc5d799c60924a36a8d34_347
a8d17c65edb97723b4ebe56373c07ee0a8b1567f_354
a016c95a288d05ea8b27c2acb3dcbe4764f2e709_229
99753a43a096b4ef9366c5e533d583784e84c3ad_158
739bd864eecb94c6de0a292846ccbaf18f4240a7_170
43fa085caa566a36e25969e2e97ffdea31eda045_129
01a8638e3f30ab7846a50539a306dc52fdf82dbd_119
2da5b3eb55017c94264e866e07818b0864a5c3c2_176
e04dc0f62a64ef583cc4dc4c706a680b2e1aa6eb_162
829d1b164c2896b8c6060a79bb6895fb586bf64f_129
844962070437fe7f89b7dc2c17e08eb06d7ed1c7_233
f90e1d3417861d7ea72b0374db53e0232b7c069e_116
0977faa48e661e7fcc5cef43d3ffa6b9d25c2452_139
5b69eb05cc2d757a032e3e8e78d0246e529ed185_198
1695c4a0bf3dbde87ef638bf0b6f4da715b657f7_105
c523a186a8e841fb4c4a521e3dbd20bd14c86c4d_107
e823fb81d263ce8786e75badb0f0f44bc52e5e6e_78
1bd194b0d0e51088ec189bae6487b604e14c0ba0_105
a8aca37ba2936ac92ae64b658a3b595eff025d36_88
756c4c1a7932c3553c365ddb070b125c1c1235d5_82
df7f3711309eca696c2529a44b78f4e6de0badb3_67
434335cd1d366058a4e591b3ff7902ba3d712f90_62
6e0e2138c79115b5b58f2a77f927f174228d7a17_69
f07d92a984474455dd525fd737a728391a2b6bb6_86
5840d9030eea063cf24dddc0d0f969885caff52e_62
22fc98e5e89a61cd4562042d0c02fe16ec2359f3_82
bd48536b13019496a7df289722c32a7199266219_107
941c0fa051a096e003c699f7747c41bf07765314_82
6868d656630344ab420014389943aece04921224_101
c71b4f6b6fea023459721b3e84a267ce1e4f3d54_55
8a9d5714f75cc03908f4a56d6294f4d7a565b8ba_77
406d953c7717cabc7d0b33cc312c35583a3f6e92_48
1a33ca03fc0df5c880a38e8955b39770bb2f86a4_54
a512cec226db446585066f21cb2ee78eb7909a02_47
f07fe2f6863eaffd02c34329d2fe3d42a6bda1f0_46
e398752440eed639b239a26a925e360b69b299f8_191
141fb7981938a04cae738acc6e1a4461178c37bf_49
717ab87a0c314e50ef5a691d5834310a412134fe_47
04acab29ba78ca3d301602c7083326ca9fb8268d_35
557eccbbd5a730515faee95c3fd93332d424bb1a_43
b50db50ae4943641b8a86e73a9dc95baa4a19980_79
bea5d30af60fb8467c4e1d6ec2491c7a027f34ae_46
de7add9cb1d2ef7f2b25e550b24a9fdee05fefb0_41
01f843ca567112849d23759e6fc29b0c6c8b7940_42
de73d3a95a35d48bee80d3e42f5f428675979f9c_37
1d0d47eb3fed4d61b55a7038a12b1a450c99d492_68
ef2d558f3c53c0960a9e2eb5f0ab267ecdaed011_53
57659cfc0f021ef7ddf42433f106c987277d2de8_38
a0e21fe41beed01ba2ae497754705b95ffdd06e1_30
31e59f4d50eaaa10d406bd23e365453ecda4af54_29
3d618eac050328b7ab982248e773a35b0833c423_27
4c47950e31746d3b14dfdc5b1a8c5812549014dd_39
e24f8a79523089a5e1478ab6d45ae30e95ae92e9_29
94d7e5203b4b5cd437c9ec87770a366c06dcfb15_42
e8bbe0dbf9a15b5ea9c741c350538a8a84b78834_27
2d07ccc4f13dfd948ef178c1b9efde6cc8c5b3fa_22
50d0c55a83c434ad490f295f387e09dfc6715e0b_31
75ff38e18e6f31328fe328f6f886a0825f11a052_22
15801d3a29f7961f425dada58e917cfddcc2ddd6_31
2a87298f5ccf55068ce47e58101460d7d6635646_23
2e394ea765bf21bafd2b1cb1ffcf23c07e345320_25
88e81cf401e23c4170e58b69bd9792a1eab54ef1_37
8fdd827e5aaebff23e7141d531d4473c85c9e4b3_24
f15da23c46bf7d4e8ed0100e0f51d32c907361ce_33
2efa5b52e5f4ad40b6eeda59eb49ca243500d8a5_23
5a6b109fba234a42b8b3dd77923a50bc85643101_22
714697f848cfe6d73032442955fca9a8b52ba7cc_51
c152628fcd8d2e43326e430ed5962bbb941ef011_21
8717bf65a60a565c009620537308e2f0e57f4a16_27
a2eb76bc4cedfebc1d711ed6d6a3fba9adf1fcc5_51
bb6c88013cfdc29d332f6e40fbbc6de3363b7ed1_23
c1819f88d8afe2c8ab4cf73a76cf7bc3651c1ba5_37
05972c797a2c6f8d1fa32d9b5f0caf88d0b15d16_17
53d48125e33c5a57830b5c928bd94c5acc0d3e0d_50
c6d2c46f3a236eb0b657717fd2cfcee1dcf063ac_19
e4143d344fd4893179ce0a22ca5bc09186a0d42e_22
6864ed4c9ab0f2a5d7a61b174eb392c47dbf7431_21
7281a300e2fbacbfb132bc33179513ae99bc9a2f_21
87a9a37400a625a85e0f1bba40d2834aa50f42e5_61
4d43cbbc7f7eda599d52b0cfb45eb9e680e5f73f_19
77d331b6c643673e276b62b2e15516237f8ccc95_20
d089ac883663426a904bc9f2931c308f192daf21_17
e15cb9898273060c229841f7fb78909e71605eda_26
052e375b1b8b03de9eabd6a56c290fd8bf2bdce8_21
3f0e54d8c01430218d451533f9f0a51a73c940c2_14
6081f75ff97eecd031bcbf61649012efaf8bdb9c_37
8aedb764b91c410e79af9a410626e2d190432588_18
a9b2531cf2d5178e21d4aaa7dcc622e7606cce16_14
05a727ecc8072d8ff4e676fe70250eeb661e2aa5_11
223ed0fe0e2696e07d5888992182926f8cdb6d2f_11
2c7f5ae8e54dc84205013a4ace2b3f486aee8903_13
4cfa51b52bf80fb0132dbeee7c77311d55ebbbc3_14
4db83ae1a5b02c7637ee985a00935ea6da0034a9_10
92e8d7273417b4283b07356893bcd9335a41966a_23
f590b48a342ddad6a9b77178cd18445c05376fd3_13
0201701ea9ed0304fa8260cdc5f8b790895c17d6_14
8c5a62d7716b4a5bfe911645a800d36e59afe6dd_18
29deeb67c772ac4044ec71744905921a9f68ee5c_9
44f11409dfc8e3a951af2ffe4c26da1f8b26ba60_26
4b8ba3c5e3cb0ebb0c0353e28b73998824a941fc_8
569d0c5bc6ed01e1b559e01eed5685ae66ab2228_12
7e42418129ea5aaea8d3149fba519f32ca6ad996_26
84e420408460fd0519b7dc846f8239a6e1aba082_9
a2357d1f3dd7275476e8e5d782deeab3038aa403_13
35ec737728ca4e6f498a12719de09b79099c27d5_12
ef02f2a97f01c3be157c52e25513b9477856f56c_6
0e12e913708a090ec2aa02d10086882f87c721b3_15
c50196f46f1ce669b41fb6cf7f2782521bf1faba_8
d112d3d56666affd015d5908211b0a54b14fe6ce_5
04ef7824b6bc77c4bd755fa360e2903b5a018a47_8
1d6727aeab4b54c40b50810aa80e17a32ef0dd19_12
2e7729e1d1a799dd75d5d1cc19cd093e6eb128a4_4
351b1077e3e4b89d87dcdfc9ec20a475ea190eb7_22
44cb591706f5d8e6a8f94fb23b187c60f93fc502_4
5eed2099d239d05fd3f578af04d0a8eb81719fe0_9
7551555993b7191609cab1b611545d5ac858283b_23
a54f54dd3bfac045239dea1ce480d26dc5beac41_6
b08bcb283d7095325bdb4399e853e6aea7bd30df_7
c6abc78693ef3bcc644f9e819d7512fbed1ab6f8_5
d5f4ab857f9cd02ba17760a079c979c29f4a826d_7
edf646d1ce9e460020299f0cce2f2371dfaf8ffc_5
027bbb34b140df15b28de6788e154eb7a3c25256_8
05eef0c777d38968907e549ac5fda0fae976569e_11
219b58b56075653d073f23e624f1231d2d51b2a9_4
25c273820cb13e883701fe04d21f30671394dfdd_4
34f4882d7a687b1704629152f3160c66a50bb507_4
375232b46264968ca639d68b54706febd5940eef_7
3c69faa7cccdb765a46ca2cc9a11335d17a37c5b_3
401b74102d854e9194b6e40deaf762a0788ba7f5_3
446be04449e7bd4dbb1332aab3d59b6a780bc37b_3
5a972385ad77370f1f3cd1533fafff808851a9c6_8
6f1a2a0b2723bb665470a0bc6b8e046171c1cc3b_3
70ccb08683d31f2554f0a6c1ebfe0f2ee3774d51_4
7c5d0fa3a51c0c33ae346fc5fbf66eead60a292a_3
88512c3691291ff5821c4b2d945b6605ab9e6b05_4
954b122fd38f22d87765e04c226a8cf21cacca82_8
b682ad855474e4a646f74f8cb5e4ec6ac2f5a7e0_3
c1948c6861b48991d26efd8e8d36a3347cc302c0_14
c29d5a6a583c4e4a62d2935b0e7f05c01fc19a27_3
cf71e0a1e86f46009c46d8805d2f6b63c84f4d74_4
f023571e62541334f53cd871436d376e43822c04_5
f6e387974264f02dd3fc45e7658376090ef0e0f1_5
12adbe05d11abbf6fe5db52cc8c215c39238898b_4
184630ce3169cedb863a2b41146a76d4184a382d_6
1d9f8af9ee85e3ca41d87fff16ba2b18a86c99ab_3
32657cf7f28fc0772b6f408eeeca3a3d0938ac7e_3
3f5916dbedace7fe186b6695c3853f5d611ff4b5_4
40d384cca0e8c2905e7b9c6100268cb4404a9be2_3
48896daee6d0282204948ba4f1eccecfd778d12e_3
555654bcf76e644fd57214a89ff851446edee952_5
73542d022ad821bebf391c891e25bb9126bc20d9_3
73daf1f9b09b68471ccc8848c3ad7ccede1b9001_4
78c59b6a3242068436465adb5fa041ea638fcae2_5
7b535fbe7351a10f07fa5aa08f0e7f70fd50a7de_3
8816255c36035eb9c73eceb33b50e546a27f46cd_4
8a7145b6d0d688f4e925c91ea792df5898efd471_3
909206944c59a4e31b48413b9a11c5c39a206bd2_3
941e7fb35a1187036e96695bb81d98f64f91fbbf_4
a3bf44780ae346ada7cd1e490c47e606a7127abf_4
b1093edf2c320761928a2656331c1cd92815e6e2_4
bfda99444b9daa2d2ab0ede9bda49c5da1ff96d2_3
c311ef5d8908092d7649a4b86ff34273105b6a6e_3
cb0e88950a2cc289413931c33489bdbbb2df962c_3
ce9456ca3f4ce42c2991fb9de99546e195028d77_3
ebdeec8e1685a0f5f47bf0781aab60d9dd0247a4_7
fada23666e1108b91be2e58113946ded4de67a67_3
07addf23dc12ac3650d6ac9a08d5a601263a2ea2_3
10e973bc2abbde367ebe10474e15c9612f44bc1f_5
1f1190ecbb7bb604bcee174d8c547bced08c564a_5
2cd9a5773ee03a3cfc94f7370bc8cf5470ef53b7_1
3a9ee8362853e6d4bf9798ad6dc28e740bd35618_3
4a2def16f7ec2c2acd8c9a6d338695cc8622f892_3
4e9180a82d2b0c7419c23600e56d28079de5c9ba_5
94bc4dff1b587b35c04e74d3f0f65453509fac59_1
97ad310530563be46378d9f479194c27c6ce10a1_1
d23d7a20f58396dcc5c4bf2f5e0fff05c9248d8a_3
